# Opportunistic pathogenicity in fungi can transcend species boundaries

**DOI:** 10.64898/2026.07.02.736111

**Authors:** David C. Rinker, Thomas J. C. Sauters, Adiyantara Gumilang, Olivia L. Riedling, Karin Steffen, Camila Figueiredo Pinzan, Thaila Reis, Patrícia Alves de Castro, Manuel Rangel-Grimaldo, Huzefa A. Raja, John G. Gibbons, Nicholas H. Oberlies, Gustavo H. Goldman, Antonis Rokas

**Author notes:** Corresponding authors: Gustavo H. Goldman, Nicholas Oberlies, Antonis Rokas.

## Abstract

Multiple fungal lineages have independently evolved the ability to opportunistically infect humans, imposing a major burden on public health. Opportunistic pathogenicity requires the confluence of pre-existing fungal traits that facilitate host colonization (*e.g*., the ability to grow at 37°C) and host immune filters that permit the survival of some colonizers (*e.g.,* inborn errors of immunity). Numerous studies have shown that fungal pathogens can exhibit extensive strain-to-strain variation in the ability to cause disease. Moreover, it is also well established that non-pathogenic fungi can occasionally cause severe infections. Together, these observations raise the question: What differentiates opportunistic fungal pathogens from non-pathogens? To address this empirically, we compared phenotypic, metabolomic, and genomic variation between *Aspergillus fumigatus*, an organism responsible for more than 300,000 infections per year, and *Aspergillus fischeri*, a close relative that is not considered clinically relevant. By examining 26 phenotypic traits across 16 representative strains of *A. fumigatus* and 16 of *A. fischeri*, we found that infection-relevant traits measured under *in vitro* monoculture conditions showed species-specific distributions, whereas traits measured under *in vitro* coculture with murine macrophages overlapped. Strikingly, strains of the two species also overlapped in their virulence profiles in an immunocompromised murine model of pulmonary aspergillosis; three strains of *A. fischeri* exhibited lethality rates of >50% while two *A. fumigatus* strains were among the least virulent of all 32 strains tested. Consistent with this overlap, we could not associate virulence variation with the presence of specific genomic elements, phenotypic traits, or secondary metabolites. Our results raise the hypothesis that opportunistic pathogenicity can extend beyond the boundaries of individual species. We propose a conceptual model in which the opportunistic pathogenic potential of any fungal strain is the product of complex interactions among numerous genomic, ecological, and host immunity factors.

## INTRODUCTION

Fungal disease is a major burden, annually contributing to infections of more than a billion people and more than a million deaths [1]. Of the more than 150,000 described fungal species [2] and the millions more of estimated species [3], about 200 are known to infect humans [4], with just a few dozen species accounting for the vast majority of serious infections [1]. Fungal pathogens of humans are scattered across the fungal tree of life, suggesting that opportunistic pathogenicity has arisen independently numerous times [5].

Remarkably, most fungi that infect humans are considered accidental or opportunistic pathogens [5]. Opportunistic fungal pathogens typically infect only immunologically vulnerable hosts and are not reliant on a host to complete their life cycle. Indeed, most infectious fungi are environmentally acquired, are not typically transmitted between hosts, and are not known to gain any fitness advantage from causing disease. Thus, the evolution of fungal pathogenic potential may be best understood as an interaction between a pathogen and a host that has emerged in the absence of coevolution. Instead, the pathogenicity of most fungi results from the convergence of two independent processes: 1) ecological fitting [6–8], by which pre-existing fungal traits coincidentally facilitate host colonization, and 2) environmental filtering [9], by which features of the host’s immune environment subsequently restrict which colonizers survive.

In the context of mammalian fungal disease, the most widely cited example of ecological fitting is thermotolerance [10], an adaptation evolved in response to warmer niches, such as decomposing compost or geothermal heat sources, that also allows a fungus to grow at the human body temperature. Other suggested examples include the ability to respond to (and, for some traits, actively modulate) hypoxia [11,12], oxidative stress [13], fluctuating CO_2_ levels [14], pH shifts [15], competition for nutrients [16], and the ability to biosynthesize bioactive molecules, such as melanin and other secondary metabolites (SMs) which protect the fungus and mediate interspecific interactions [17,18].

In contrast, environmental filtering occurs when the human host environment limits which fungi can grow in it. Environmental filters include variability in host immune competency due to extrinsic factors like iatrogenic immunosuppression (*e.g.,* chemotherapy-induced neutropenia [19]) or genetic factors like inborn errors of immunity [18,20]. One notable example is *Cryptococcus neoformans* infection, whose incidence dramatically increased during the HIV/AIDS pandemic [21]. *C. neoformans* disproportionately affects patients living with HIV because the virus destroys the same immune cells that combat yeast infections [22].

In this framework, the pathogenic potential of a fungus is determined by its set of traits that ecologically fit life inside a human host—sometimes described as the “cards of virulence” model [23]—as well as by the specific environmental context of human host availability and susceptibility. Jointly, these form the major components of the “damage-response” framework of pathogenesis, with virulence then being an emergent property of this complex and dynamic network of host-fungal interactions [24,25]. There are two evolutionary implications of this framework. First, there are likely many combinations of fungal traits and host environments resulting in virulence, and hence many trajectories by which a species can become pathogenic. Second, no single fungal trait or host environment is likely to fully explain the pathogenic potential of a given microbe [5,23].

Infections caused by members of the fungal genus *Aspergillus* account for millions of disease cases per year [1,26]. Approximately 70% of aspergillosis infections are attributed to *A. fumigatus* [27], a globally distributed saprophyte that the World Health Organization recently ranked among the four most critical fungal pathogens [28]. *A. fumigatus* is one of the ∼60 species contained within the taxonomic section *Fumigati* [29]. While many *Fumigati* species have been identified in clinical contexts, only a few are considered “clinically relevant” (i.e., routinely described in clinical infections but rarely exhibiting a high prevalence) [29,30]. And, like pathogenic fungi in general, those clinically relevant species are dispersed across the phylogeny of section *Fumigati*, a distribution suggesting that each species independently evolved its pathogenicity [5]. Indeed, despite *A. fumigatus* cases overshadowing those of all other pathogenic *Fumigati* species by two orders of magnitude, the two species most closely related to *A. fumigatus*, namely *A. fischeri* and *A. oerlinghausenensis*, are not considered to be pathogenic [31–33], even though *A. fischeri* is occasionally detected as the causative agent of aspergillosis (in small numbers) in various studies.

Given their close phylogenetic relationship, *A. fumigatus, A. fischeri*, and *A. oerlinghausenensis* might be expected to share similar genome sizes, gene contents, and evolutionary rates. This is broadly true; for example, virulence-associated genes are widely shared across species in section *Fumigati* [34]. However, certain aspects of the evolution of *A. fumigatus* are notably distinct from those of its close relatives. For example, phylogenetic analyses consistently show *A. fumigatus* protein-coding genes [31] and non-coding regions [35] to be evolving much more rapidly compared to its sister taxa while divergent evolutionary rates have been identified as one of the genomic factors that seems to distinguish pathogens from nonpathogens within section *Fumigati* [34]. Differences in biosynthetic gene clusters (BGCs) and their associated SMs have also been identified as being divergent between *A. fumigatus* and *A. fischeri* [33]. The *A. fumigatus* reference genomes are also consistently about 10% smaller than *A. fischeri* reference genomes and show a proportionate reduction in overall gene content [32].

A newly appreciated source of comparative information for identifying factors related to pathogenic potential is variation between strains of the same species. Early comparisons between environmental and clinical strains of *A. fumigatus* revealed that clinical strains showed greater fitness under low oxygen conditions [37]. A pangenome analysis of over 300 *A. fumigatus* strains found that clinical strains differed from environmental ones by their enrichment in accessory genes [38]. Subsequent whole genome analyses showed that *Starship* elements are a primary driver of strain heterogeneity within *A. fumigatus* [39]. Examination of the genomic and phenotypic diversity of *A. fischeri* further revealed that species considered to be non-pathogenic can also harbor substantial variation in pathogenic potential across strains [32], a finding consistent with its infrequent detection in clinical settings.

If fungal pathogens can exhibit extensive strain-to-strain variation in the ability to cause disease and non-pathogenic fungi can occasionally cause severe infections, what differentiates opportunistic fungal pathogens from non-pathogens? We hypothesized that if strain-level variation in pathogenic potential is sufficiently broad, the virulence distributions of a so-called ‘pathogenic’ species and a ‘non-pathogenic’ one will overlap. To test this directly under matched, host-relevant conditions, we comprehensively profiled the genetic, chemical, and phenotypic traits of 16 representative strains of *A. fumigatus* alongside 16 strains of *A. fischeri.* We evaluated these strains for previously characterized infection-relevant genes, pathways, and traits (*e.g.,* morphological measures, gene content, BGCs, SMs) and measured each strain’s pathogenic potential using both *in vitro* assays of mammalian macrophage responses to spores, as well as an *in vivo* murine model of pulmonary aspergillosis. We found that many infection-relevant genetic, metabolomic, and phenotypic traits readily distinguished the two species. Surprisingly, strains of the two species showed substantial overlap under *in vitro* coculture with murine macrophages. Furthermore, while *A. fumigatus* strains were typically more virulent, several *A. fischeri* strains were more virulent than the least virulent *A. fumigatus* strains in an immunocompromised murine model of pulmonary aspergillosis. The substantial overlap in the pathogenic potential of the two species provides evidence that the ability of fungi to establish opportunistic infections in human hosts can transcend species boundaries.

## RESULTS

### Phylogenomic analysis reveals that *A. fumigatus* and *A. fischeri* are reciprocally monophyletic

To characterize strain heterogeneity within *Aspergillus fumigatus*, we first retrieved genome sequence data from hundreds of clinical and environmental strains deposited in the NCBI Sequence Read Archive. We inferred the population structure of these strains, identifying seven distinct groups (Figure S1). We then selected 14 representative strains, where at least one strain was from each of the seven population groups we identified (Methods; Figure S1). For these strains, we generated long genomic reads (ONT) to augment the publicly available Illumina reads, allowing us to *de novo* assemble near-chromosome length scaffolds for each strain, which we then individually annotated. Each of the annotated assemblies contained a nearly complete complement of conserved, single copy genes (BUSCO completeness of >99% for the Eurotiomycetes dataset). The quality of the *A. fumigatus* genome assemblies and annotations mirrored the quality of the results we previously reported in 16 strains of *A. fischeri* [32], and allows for informative comparisons between these two species. These 14 strains plus the two common reference strains Af293 and CEA10 comprised the 16 *A. fumigatus* focal strains of our analysis (Table 1). We also assembled and annotated the genomes of two more distantly related species from section *Fumigati*, namely *A. lentulus* and *A. fumisynnematus*, which we used as outgroups.

**Table 1.**
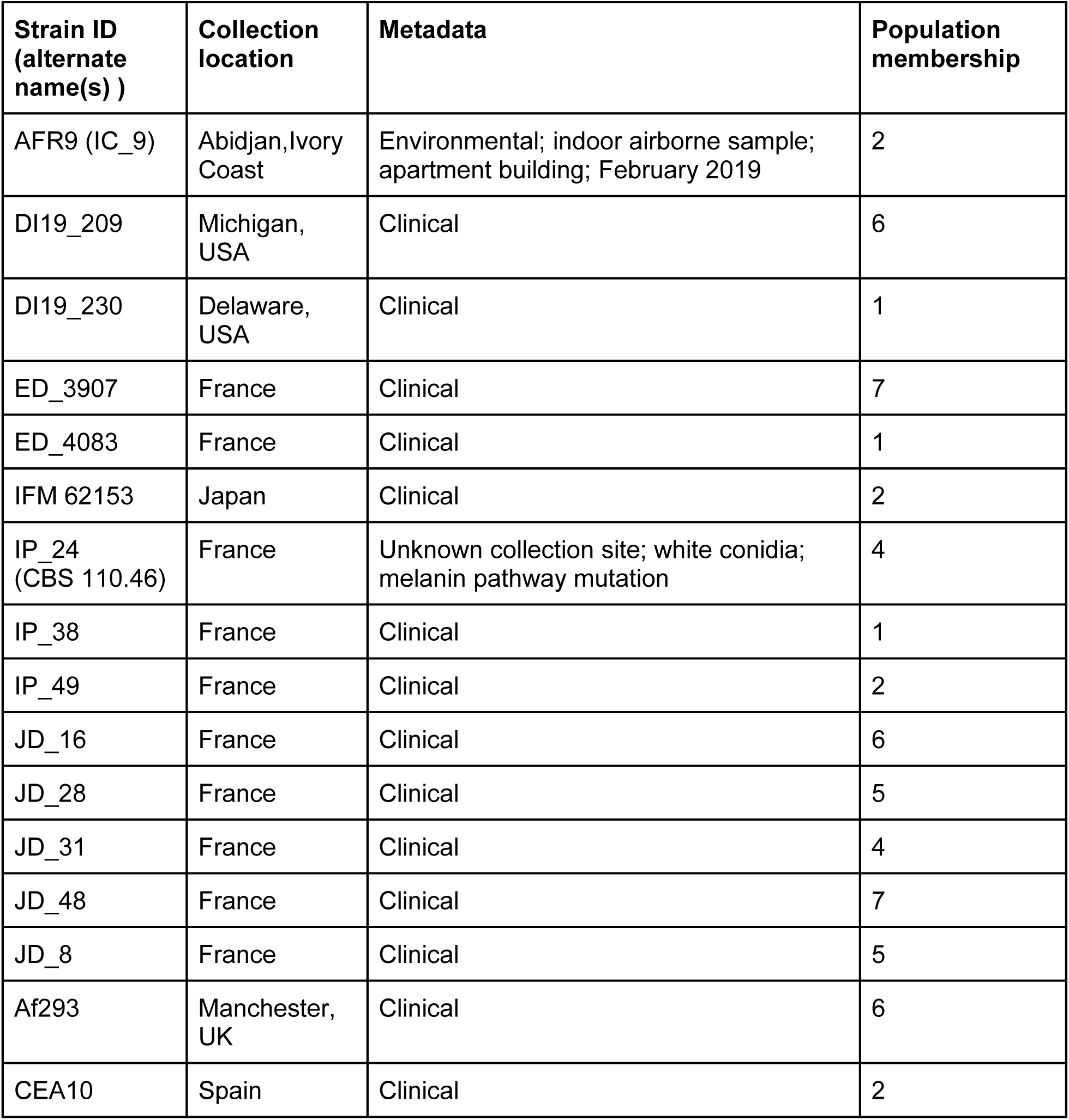
*Aspergillus fumigatus* strains used in this study.

| **Strain ID**<br>**(alternate**<br>**name(s) )** | **Collection**<br>**location** | **Metadata** | **Population**<br>**membership** |
| --- | --- | --- | --- |
| AFR9 (IC_9) | Abidjan, Ivory<br>Coast | Environmental; indoor airborne sample;<br>apartment building; February 2019 | 2 |
| DI19_209 | Michigan,<br>USA | Clinical | 6 |
| DI19_230 | Delaware,<br>USA | Clinical | 1 |
| ED_3907 | France | Clinical | 7 |
| ED_4083 | France | Clinical | 1 |
| IFM 62153 | Japan | Clinical | 2 |
| IP_24<br>(CBS 110.46) | France | Unknown collection site; white conidia;<br>melanin pathway mutation | 4 |
| IP_38 | France | Clinical | 1 |
| IP_49 | France | Clinical | 2 |
| JD_16 | France | Clinical | 6 |
| JD_28 | France | Clinical | 5 |
| JD_31 | France | Clinical | 4 |
| JD_48 | France | Clinical | 7 |
| JD_8 | France | Clinical | 5 |
| Af293 | Manchester,<br>UK | Clinical | 6 |
| CEA10 | Spain | Clinical | 2 |

To evaluate per-strain genome content, we consolidated genes from all 16 strains of *A. fumigatus* into a pangenome of 14,444 single-copy pangenes (Methods). Of the pangenes, 8,429 (58%) are present in every strain (core genes), with the remaining 6,015 genes present in only some strains (3,774 accessory genes) or just one strain (2,241 singleton genes). Overall, the pangenome of the 16 *A. fumigatus* strains was smaller than that of a comparable 16-strain *A. fischeri* pangenome (16,148 single copy; 9,053 core; 7,095 accessory or singleton) and is consistent with the previously described gene content both within and between these species [32,38]. The accessory gene component of the *A. fumigatus* pangenome monotonically increased without converging (Figure S2; Heap’s law γ = 0.854); this result is similar to previous analyses of the *A. fumigatus* pangenome [38,40,41] and what we observed in the *A. fischeri* pangenome [32], suggesting that the sampled pangenomes of both species are open.

To reconstruct evolutionary relationships, we inferred a strain phylogeny using orthologous, single copy Eurotiomycetes genes (BUSCOs) present in the 16 *A. fumigatus* strains, the 16 *A. fischeri* strains, and the genomes of *A. lentulus* and *A. fumisynnematus*, our two outgroup species (Figure S3). Strains of *A. fischeri* and of *A. fumigatus* were reciprocally monophyletic on the inferred phylogeny. Within each species, genetic distances were similarly distributed with only slightly higher rates of evolution among *A. fumigatus* strains than among *A. fischeri* strains (median patristic distance = 0.0014 and 0.00083 nucleotide substitutions/site, respectively). The observed genetic distances within each species dwarfed genetic distances between the two species, with a median pairwise distance between *A. fumigatus* and *A. fischeri* equal to 0.088 nucleotide substitutions/site, a value 80 times higher than the median of all intraspecific genetic distance for both species (0.0011 nucleotide substitutions/site).

Within *A. fumigatus*, two strains (JD48 and ED3907) were substantially diverged, forming a sister clade to the other strains; these two strains belong to a distinct subpopulation of *A. fumigatus* (Population 7, Figure S1) and exhibit the highest pairwise distance (6.0 × 10^−3^ nucleotide substitutions/site, almost three times higher than that between any other intraspecific pair) among our focal 32 *A. fischeri* and *A. fumigatus* strains. Excluding strains JD48 and ED3907, the highest pairwise distance among the remaining strains of *A. fumigatus* was between the two strains from Population 1 (IP38 and ED4083; nd Af293 from Population 6 (2.3 × 10^−3^ nucleotide substitutions/site); this distance nearly equaled the maximum divergence seen among the *A. fischeri* strains (2.2 × 10^−3^ nucleotide substitutions/site between CBS 150754 and CBS 147335).

### The distributions of *A. fumigatus* and *A. fischeri* phenotypic traits measured via *in vitro* monoculture assays rarely overlap

To gain insights into levels of strain heterogeneity (intraspecific polymorphism) and interspecific divergence, we individually assayed *in vitro* monocultures of each of the 32 strains for 21 quantitative infection-relevant traits associated with the immunogenic composition of asexual spores (conidia), the ability to form biofilms, and radial growth rates under a variety of abiotic stressors (Figure 1A).

**Figure 1.**
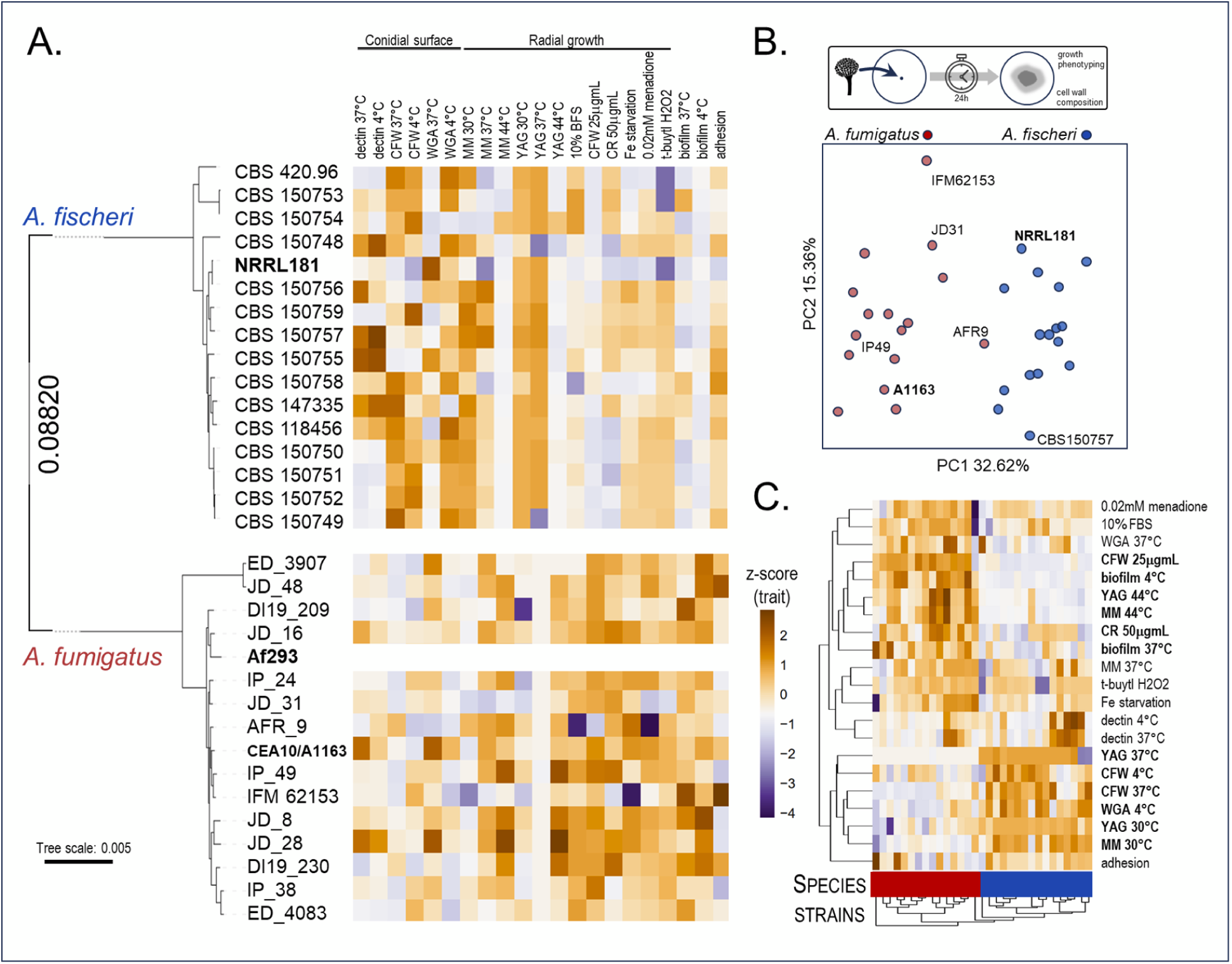
The phenotypic profiles of *A. fumigatus* and *A. fischeri* inferred from monoculture assays do not overlap. **A.** Phylogenetic tree of the 16 *A. fumigatus* and the 16 *A. fischeri* strains (left) along with their values for 21 phenotypic traits measured using *in vitro* monoculture assays (right). The phylogeny was inferred from 3,100 single copy BUSCO genes. Strains in bold are commonly used reference strains. The heatmap represents z-scores for each of the assayed traits; z-scores are calculated across all strains. Abbreviations: WGA (wheat germ agglutinin); CFW (calcofluor white); CR (Congo red); MM (minimal media); YAG (Yeast Extract-Glucose); t-butyl (tert-butyl hydroperoxide); fetal bovine serum (FBS). **B.** PCA of 21 traits across 32 strains of *A. fumigatus* and *A. fischeri.* **C.** Traits clustered by z-scores; traits that were significantly different between the two species are in **bold** (see Table S2).

Overall, the variances within individual traits among *A. fischeri* and *A. fumigatus* strains revealed by the principal components analysis showed clear, species-specific differences, with just one strain of *A. fumigatus* (AFR9) appearing to bridge interspecific phenotypic space (Figure 1B). The divergence between the two species was largely driven by differences in radial growth at 44°C, the ability to form biofilms at 37°C, and by variation in levels of the immunogenic surface polysaccharide chitin (CFW) (Figure 1C; Figure S4). Generally, *A. fumigatus* strains exhibited some infection-relevant traits—such as early immune evasion, thermal tolerance, ability to grow in low-nutrient conditions, and biofilm formation—with greater magnitude and higher frequency. However, even these traits were rarely entirely species-specific and occasionally also reached high magnitude in *A. fischeri*. Other traits related to levels of such cell wall carbohydrates as beta-glucans (dectin) and amino-sugars (WGA) appeared variable both within and between species (Figure 1A). Indeed, proxies for beta-glucans (dectin and CA) strongly drove intraspecific variation within both species, followed by traits reflecting susceptibility to oxidative stress (t-butyl H_2_O_2_) (Figure 1C).

To measure levels of intraspecific and interspecific phenotypic variation, we inferred a phenetic tree from these traits (Figure S5). The observed phenetic distance between the two species (median = 7.2, s.d. = 1.06) was slightly higher than the intraspecific distances in either *A. fumigatus* (median = 5.0, s.d. = 1.78) or *A. fischeri* (median = 5.2, s.d.= 1.36). The small difference in intraspecific and interspecific levels of phenotypic variation does not reflect the much larger degree of genetic variation (median interspecific genetic distance = 0.09, s.d.= 5.7e-4, which is 70- and 108-times that of the intraspecific genetic distances in *A. fumigatus* and *A. fischeri*, respectively).

To better understand the relationship between genetic and phenotypic variation, we next compared the phenetic tree of traits to the molecular phylogeny (Figure S5). The two trees were moderately correlated (Spearman rho = 0.603) and exhibited a quartet distance [42] of 0.318 (i.e. less than one-third of all quartets of strains differed topologically between the trees; Figure S5). In contrast, quartet distances between the phenetic and the phylogenetic tree within each species were approximately twice as high (0.634 and 0.535 for *A. fumigatus* and *A. fischeri*, respectively) as the quartet distance between species.

Taken together, these results reinforce the taxonomic consensus that these two species are phenotypically and genetically distinct [43]. However, our data also reveal substantial intraspecific phenotypic variability within *A. fumigatus*, illustrating the importance of considering strain heterogeneity when examining variation within and between fungal species.

### Few virulence-associated genes are specific to *A. fumigatus*

Perhaps the most noteworthy species-specific difference between *A. fumigatus* and *A. fischeri* is their vast difference in prevalence in clinical cases of aspergillosis [30,44]. Given the high number of shared orthologs between these two species, we compared the genomes of all strains for the presence of 207 genes previously described as having prior association with virulence in *A. fumigatus* [32,33].

No single strain of either species contained the full complement of all 207 genes. Unsurprisingly, the *A. fumigatus* reference strains Af293 and CEA10 contained nearly complete complements (206 and 205 genes, respectively). Nearly all other strains of *A. fumigatus* showed similar profiles with very few and sporadic absences; only two *A. fumigatus* strains lacked more than three genes with the lowest count being 201 in DI19_230 (Figure 2). The gene *chsG* (AFU3G14420, a putative class III chitin synthase (Mellado et al. 1996) was notable for being present only in the Af293 strain. Comparison of gene content between both species via reciprocal BLAST searches of the respective pangenomes showed that *A. fischeri* strains also contained most virulence-associated genes (97%), with just five genes appearing consistently absent: *chsG*, *fmaA* (*fma-TC*; AFU8G00520), *fmaB* (AFU8G00370), *fmaG* (AFU8G00510), and *pes3* (AFU5G12730).

**Figure 2.**
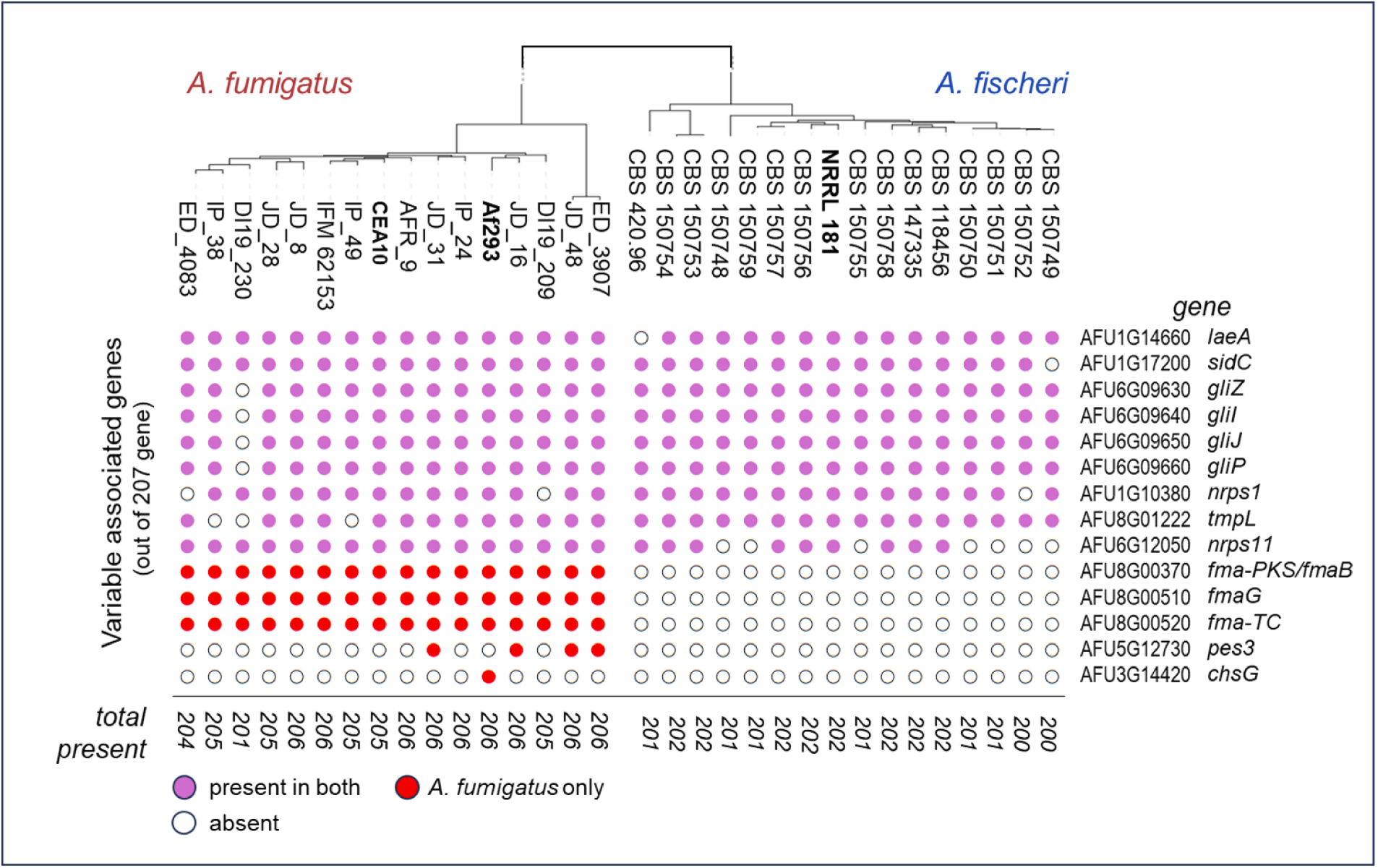
Only 14 of the 207 genes previously associated with A. fumigatus virulence are variable within and between species. There are 207 previously characterized virulence-associated genes in *A. fumigatus*. Of these, 193 are present across all strains of both species and are not shown here. What is shown is the variation in the presence and absence of the remaining 14 variably present, virulence-associated genes (one per row). Gene presence is denoted by filled circles; purple circles correspond to genes found in both species and red circles to genes found only in *A. fumigatus*. Columns correspond to the 32 strains of *A. fumigatus* and *A. fischeri* examined. The bottom row shows the total number of virulence-associated genes present in each strain.

Of these five genes, *fma-TC*, *fmaB*, and *fmaG* are part of the biosynthetic gene cluster (BGC that produces fumagillin, an SM with known antimicrobial activity [45] and a cluster that has previously been reported as partially or wholly missing in some *A. fumigatus* strains [46]. In a host, fumagillin can potentiate host immune evasion [47] and effect tissue damage [48], attributes that could promote infections and virulence; it can also exhibit antiangiogenic properties [49]. The final gene*, pes3*, is notable because its disruption has been shown to have positive impact on virulence [50], yet it is absent from all *A. fischeri* strains and is the most frequently absent virulence-associated gene in *A. fumigatus* (present in just 4 of the 16 strains). The nearly universal conservation of virulence-associated genes in *A. fumigatus* and *A. fischeri* strongly suggests that the variation in the clinical relevance of the two species cannot be solely attributed to the presence of any specific gene previously associated with *A. fumigatus* virulence.

### BGCs are more variable within *A. fumigatus* than within *A. fischeri*

BGCs are genomic features that are responsible for producing a variety of SMs. Typically, SMs are non-essential, conditionally biosynthesized compounds that benefit the fungus, primarily by augmenting or ameliorating local environmental conditions to promote growth and for mediating ecological interactions with other organisms [51]. SMs are thought to play key roles in *A. fumigatus* pathogenicity [52]. Some fungal SMs (*e.g.,* gliotoxin) have even been detected in the blood of aspergillosis patients, meaning their effects may be not only local but also systemic [53–56]. Many of the 207 virulence-associated genes are located within BGCs and are associated with the production of SMs implicated in virulence. To identify likely BGCs in our strains, we relied on both the homology and synteny of constituent genes to identify BGCs in each of our strains (Methods).

Despite *A. fischeri* having a larger total BGC count compared to *A. fumigatus* (39 vs. 33 BGCs appearing ≥ 90% complete), it had slightly fewer complete BGCs (23 vs. 26 BGCs appearing 100% complete). Of the 32 complete BGCs, 19 were shared while 5 were unique to *A. fischeri* and 8 unique to *A. fumigatus*. Conservation of BGC content was higher within *A. fischeri*: 87% of its complete BGCs appeared in all 16 strains, whereas only 65% were conserved in all 16 *A. fumigatus* strains. Furthermore, *A. fischeri* harbored more incomplete BGCs (16 vs. 6), while *A. fumigatus* had more fully absent clusters (14 vs. 8). Overall, the two species share substantial biosynthetic potential but can be distinguished both by their BGC repertoire and the degree to which it varies among strains of the same species (Figure 3B).

**Figure 3.**
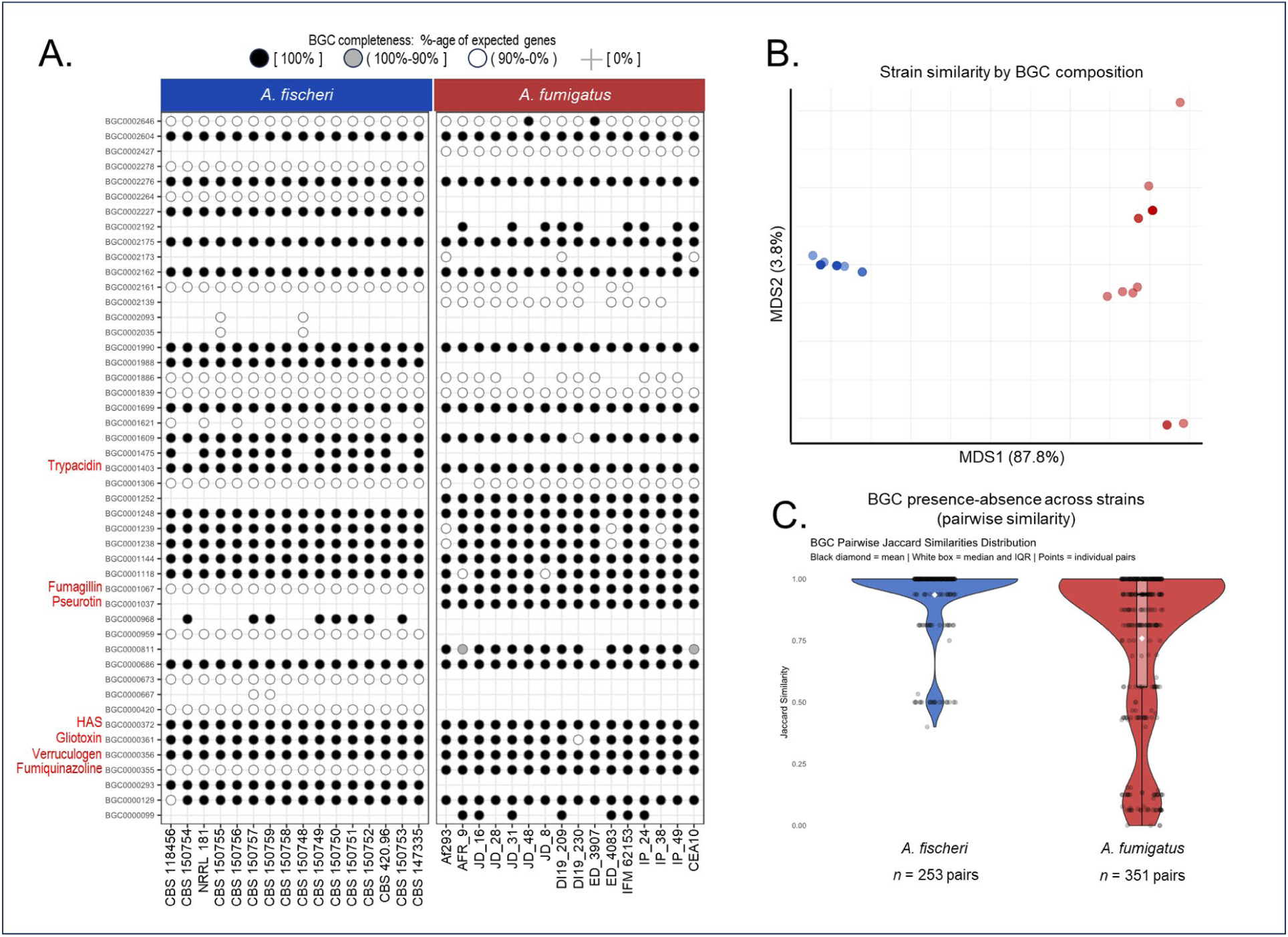
Most BGCs are shared between species but are more variable within *A. fumigatus*. **A**. Distribution of the presence/absence across strains of all cataloged BGCs that were detected in at least one strain of either species. **B**. Pairwise multidimensional scaling (MDS) analysis of BGC presence between all strains. **C**. BGC variability among strains illustrated by pairwise Jaccard similarity of the presence-absence profiles of all intact BGCs (≥ 90% complete) among strains of each species. BGC similarity distributions differ significantly between species (KS test and Mann-Whitney U test: both p < 2.2e-16).

BGCs are among the most variable features of fungal genomes and tend to be lineage-restricted [57,58]. Using pairwise Jaccard distances of BGC presence–absence profiles across all strains, we found that BGC content was significantly more conserved among *A. fischeri* strains than *A. fumigatus* (Figure 3C; 0.20 difference in median, Mann-Whitney U: p < 2.2e-16).

Interestingly, several BGCs were mutually exclusive among *A. fumigatus* isolates (Jaccard similarity = 0). For example, the monascorubrin BGC (BGC0000099) occurred in 7 strains, none of which carried hancockinone A (BGC0002646) or fumihopaside A (BGC0002173), and vice versa. The FR901512 cluster (BGC0002192) also never co-occurred with BGC0002646. Notably, such mutually exclusive BGCs only occurred in *A. fumigatus*, and encompassed both single-gene polyketide synthase clusters (BGC0002192 and BGC0000099) as well as larger clusters as illustrated by the six-gene BGC0002646 that was complete only in the two most diverged strains, JD_48 and ED_3907 (Figure 3A). Assuming that BGCs present in both species were also present in their common ancestor, their higher level of variation within the *A. fumigatus* branch suggests that they are evolving more rapidly in *A. fumigatus*. This pronounced intraspecific variability in *A. fumigatus* compared to *A. fischeri* is consistent with earlier work (Lind et al. 2017) and suggests that BGC variability is taxon-specific and may represent an evolutionary route to SM specialization.

Six BGCs are responsible for the biosynthesis of seven SMs that have been implicated as virulence factors in *A. fumigatus*: gliotoxin (BGC0000361), fumitremorgin A/B and verruculogen (BGC0000356), fumiquinazoline (BGC0000355), trypacidin (BGC0001403), pseurotin (BGC0001037), and fumagillin (BGC0001067). Except for the gliotoxin BGC, which was absent from strain DI19-230, all were complete and present across *A. fumigatus* strains. In *A. fischeri*, only the fumagillin, fumiquinazoline, and pseurotin BGCs were partially or completely absent while BGCs for fumitremorgin/verruculogen, gliotoxin, and trypacidin were complete in all strains [32].

Overall, virulence-associated BGCs were typically conserved across all strains of both species. However, compared to *A. fischeri, A. fumigatus* showed higher intraspecific BGC variability, including strain heterogeneity in the presence of a well-known BGC implicated in virulence (gliotoxin), mutually exclusive clusters, and more rapid evolutionary turnover. Such taxon-specific differences in BGC dynamics raise the hypothesis that *A. fumigatus* exhibits greater heterogeneity in its SM biosynthetic potential.

### *A. fumigatus* and *A. fischeri* metabolites form species-specific chemical spaces

The metabolites produced by *Aspergillus* range beyond SMs paired with known BGCs, and include several whose biosynthetic pathways have yet to be identified. To characterize the metabolic diversity within and between *A. fumigatus* and *A. fischeri*, we assayed extracts from cultures of each strain grown at both 30°C and 37°C using a metabolite identification protocol (i.e., dereplication) via ultraperformance liquid chromatography–mass spectrometry [59,60] (Methods).

We identified 55 metabolites among our 16 *A. fumigatus* strains, a quantity comparable to the 62 we previously identified from the 16 *A. fischeri* strains [32]. Strikingly, only 11 metabolites were shared between the two species (Figure S6). Moreover, unlike in *A. fischeri* where certain compounds were exclusive to 37°C, none of the compounds detected across the *A. fumigatus* isolates were specific to this infection-relevant temperature (Table S1).

We next assessed the chemical diversity of all metabolites across strains and treatments by encoding each compound’s structure as a chemical fingerprint. Overall, the two species produced markedly distinct metabolite structural profiles, which cleanly separated them without prior labeling (Figure 4A). In contrast, metabolite profiles varied less among individual strains within each species. Temperature-dependent shifts were markedly more pronounced in *A. fischeri* than in *A. fumigatus* (Figure 4A, B); while *A. fischeri* strains exhibited considerable variability between 30°C and 37°C, nearly all *A. fumigatus* strains showed minimal shifts, with their chemical space at 37°C closely mirroring that at 30°C (Figure 4C; Figure S7)

**Figure 4.**
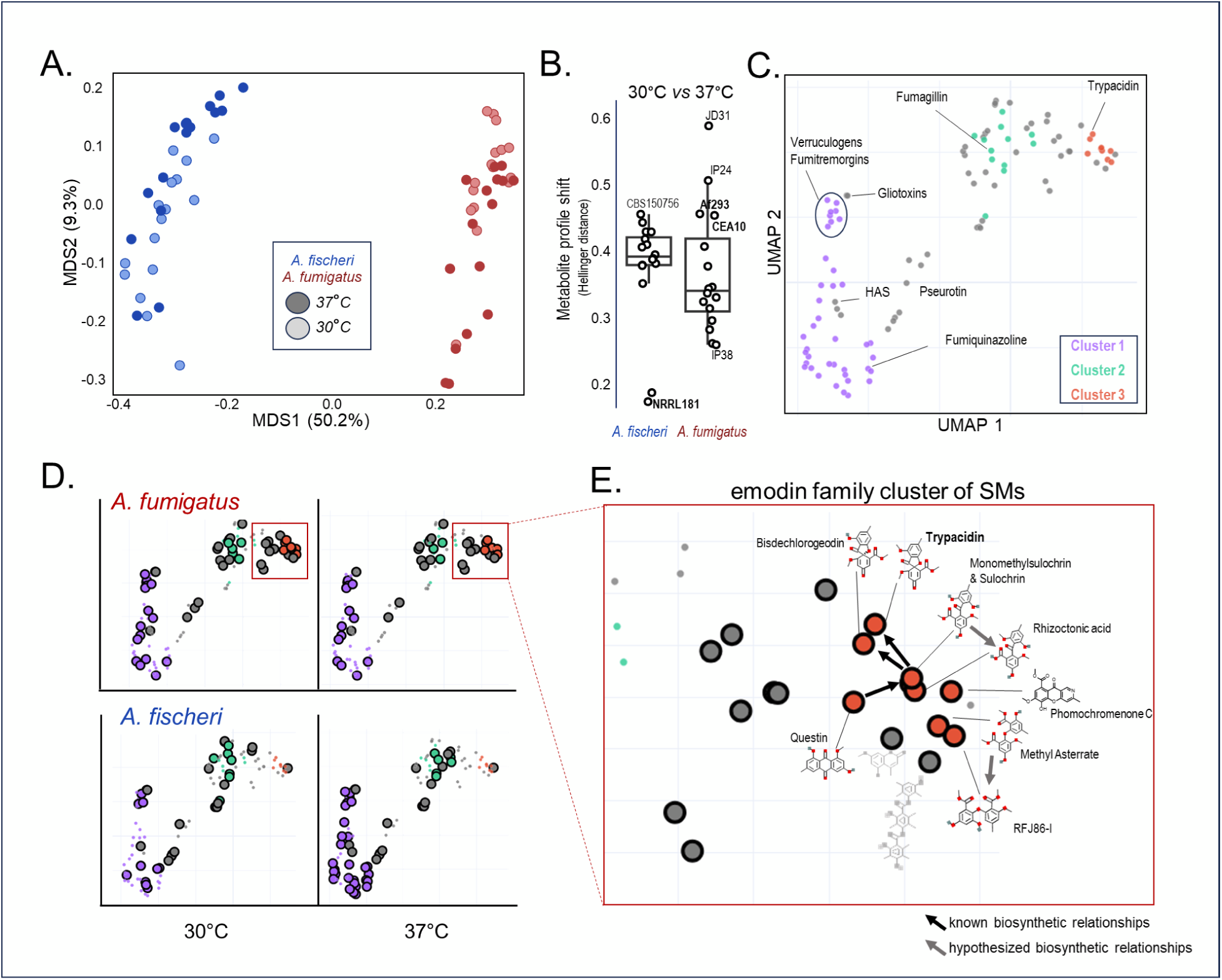
Metabolite diversity strongly distinguishes the two species. **A.** Chemical structure composition of all detected metabolites in each strain and treatment. Multidimensional scaling (metric MDS) of strain- and treatment-level chemical structure profiles derived from summed Morgan fingerprints (count-based). Pairwise distances were calculated using the Hellinger distance. Points are colored by species (*A. fumigatus* in red, *A. fischeri* in blue) and filled according to treatment temperature. Axes labels indicate the percentage of total variance explained. **B.** Temperature-induced shift in chemical structure composition for each strain. Hellinger distances between the (summed Morgan) fingerprint profiles of each strain grown at 30 °C and 37 °C, separated by species. Points represent individual strains; reference strains and max-/min-shifted strains are labeled. **C.** Chemical space of all detected metabolites. UMAP embedding (*n*-neighbors = 15) of 109 individual metabolites, constructed from Hellinger distances between their counted Morgan fingerprints. The position of each point reflects structural similarity, revealing the overall chemical diversity of the detected metabolite repertoire. HDBSCAN clustering (minimum cluster size = 6) identifies three structurally coherent groups or clusters (colored); grey points represent metabolites not assigned to any cluster. Selected, virulence-associated SMs are labeled. **D.** Treatment- and species-specific chemical spaces. The same UMAP embedding of all 109 metabolites as in the previous panel, now separated into each of four species–temperature combinations. Bolded points indicate metabolites present in that specific combination. **E.** Region of chemical space that appears exclusive to *A. fumigatus.* Detail of the emodin family with many metabolic precursors and byproducts in evidence. A selection of these compounds and their biosynthetic relationships are shown.

We then used these chemical fingerprints to construct a chemical space based upon the chemical structures of all detected metabolites in both species (109 compounds), and then clustered them based upon structural similarity. We found that more than half of the compounds (62) fell into one of three structural clusters, while the remaining compounds (47) were more structurally distinct (Figure 4C). SMs associated with seven of the virulence-associated BGCs discussed above (Figure 3A) were found equally within- and independent-of, structurally similar compounds (Figure 4C). Several clustered chemical structures reflected known biosynthetic relationships. For example, verruculogens and fumitremorgins, which are synthesized by the same BGC [61,62], comprise a distinct structural group in our inferred chemical space. Similarly, gliotoxin and bis(methylthio)gliotoxin, two metabolites associated with the gliotoxin biosynthetic pathway [63], appeared as nearly overlapping points in chemical space.

The metabolites present at both temperature conditions were very structurally similar within the species; even between species, similar structural spaces were covered despite the fact that many of the individual compounds were different (Figure 4D). The only notable difference were the eight compounds comprising Cluster 3, along with 10 adjacent compounds, all of which were wholly exclusive to *A. fumigatus* and detectable under both temperature conditions. The majority of these 18 compounds contained members of the emodin family of fungal natural products [64]. Emodins are all structurally similar (Figure 4E, S11), and biosynthetically interrelated through branching pathways involving multiple BGCs across fungi [64].

Trypacidin is a cytotoxic SM associated with *Aspergillus* spores and early stages of infection [65,66]. Among our strains, trypacidin was detected in 12 strains at 30°C and in 9 strains at 37 °C. Interestingly, neither of the *A. fumigatus* reference strains (Af293 and CEA10) showed trypacidin presence at either temperature despite the fact that this compound has been previously reported in Af293 [66,67]; trypacidin’s absence in CEA10 was less surprising given that it has not been reported previously and that a key gene in the BGC may be broken due to a frameshift mutation [68]. This variable presence is consistent with previous work showing that trypacidin production in *A. fumigatus* can be both temperature- [69] and media-dependent [70]. Notably, the BGC associated with trypacidin biosynthesis (BGC0001403; Figure 3A) was present and complete across all strains of both *A. fumigatus* and *A. fischeri*, yet this SM was not detected in the latter species [32], suggesting that conditions for its activation may vary between the two species.

### In *vitro* cocultures with murine macrophages reveal pervasive phenotypic overlap between *A. fumigatus* and *A. fischeri*

Invasive aspergillosis typically begins when airborne spores access susceptible host tissues. In pulmonary aspergillosis, spores initially deposit on alveolar surfaces of the lung. There, the spores encounter host macrophages which recognize and engulf them; these macrophages also may initiate an inflammatory response to recruit additional host defenses. Only spores that survive this early immune response germinate and invade the lung epithelium as hyphae; macrophage evasion therefore constitutes the critical first hurdle to invasive disease.

To further explore how strain heterogeneity might influence pathogenic potential, we modeled this early host response *in vitro* by evaluating how conidia from each strain interacted with murine macrophages in two ways (Methods): spore killing by macrophages, and cytokine signaling by macrophages in response to fungal spores. First, we simulated the early encounter by measuring macrophage clearance of conidia over 24 h and then scaled the result to the highly virulent clinical strain A1163 (a derivative of CEA10) (Figure 5A). Conidial susceptibility to macrophage killing varied widely, with *A. fumigatus* strains occupying both extremes. At the high end of survival, the spores of five *A. fumigatus* strains showed similar robustness to macrophages, which aligns well with the species’ designation as a pathogen. Notably, one *A. fischeri* strain (CBS 150758) also showed limited susceptibility and was statistically indistinguishable from the *A. fumigatus* strains (unpaired two-sample t-test, Bonferroni-corrected p > 0.05). At the low end, five more *A. fumigatus* strains appeared most susceptible to macrophage killing, showing lower robustness than all the “non-pathogenic” *A. fischeri* strains.

**Figure 5.**
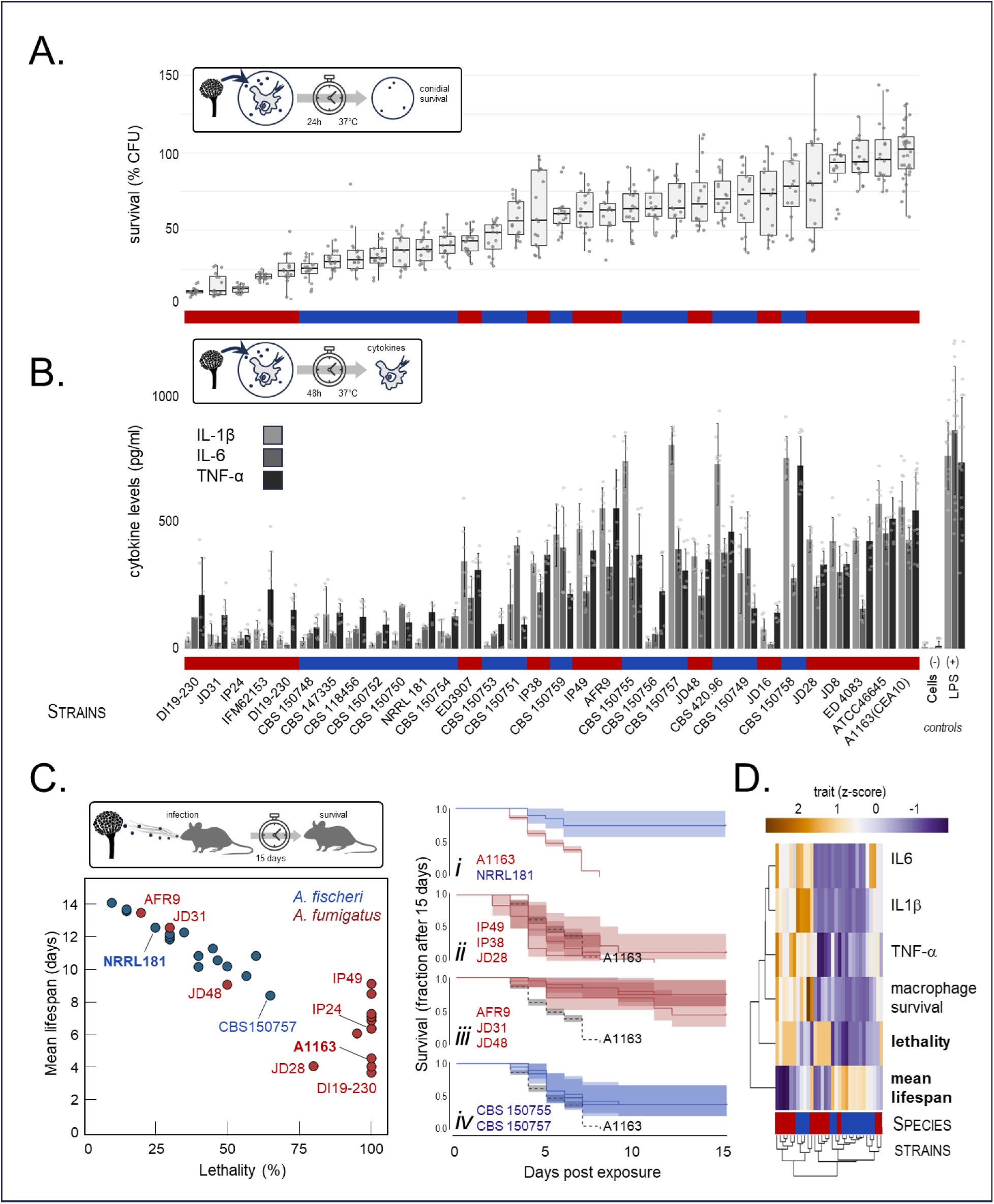
*A. fumigatus* and *A. fischeri* strains overlap in both early immune response and pathogenicity. **A**. Survival of conidia (asexual spores) from all strains as estimated by the number of colony-forming units (CFU) recovered after 24 h of exposure to bone marrow-derived macrophages (BMDMs) at 37 °C. Points represent individual assays normalized to the mean spore survival of A1163 (a derivative of CEA10). Boxplots show the median and interquartile range; whiskers show max/min values of data that are within ±1.5× the interquartile range, respectively. Strains are ordered (left-to-right) by their median conidial survival (low to high). **B.** Pro-inflammatory cytokine levels produced by BMDMs after 48 h incubation with conidia at 37 °C. Lipopolysaccharide (LPS) and cell medium were used as positive and negative controls, respectively. Bars represent the mean of eight separate measurements carried out across two replicates. Error bars show standard deviation. **C.** Virulence profiles of *Aspergillus* strains in a murine model of pulmonary aspergillosis. For each strain, 20 immunosuppressed mice (two cohorts of 10) were exposed to spores, intranasally. *Left*: Summary of survival outcomes at day 15. Each point represents the pooled cohorts of all mice exposed to one strain. Points report the overall lethality (x-axis) and the mean lifespan (y-axis) of all mice exposed to each strain. Points are colored by species and selected strains discussed in the text are labeled. *Right*: Selected Kaplan-Meier survival curves (percent survival vs. days post-infection). Subpanels show: (i) reference strains of each species, (ii) the three most virulent strains overall, (iii) the three least virulent *A. fumigatus* strains, and (iv) the two most virulent *A. fischeri* strains. Shaded bands represent pointwise 95% confidence intervals for each survival curve (log-transformed, Greenwood method). PBS-treated negative controls had 100% survival through day 15. **D.** Heatmap of z-scores for all *in vitro* and *in vivo* phenotypic traits. Traits and strains are hierarchically clustered by trait (vertical) and by strain (horizontal). In panels A, B, and D a colored annotation bar along the x-axis reports species membership of the strains (red for *A. fumigatus* and blue for *A. fischeri* strains).

The early host immune response typically features the release of proinflammatory cytokines at the site of infection [71,72]. To address this, our second assay measured the levels of three key proinflammatory cytokines (IL-1β, IL-6, and TNF-α) secreted by macrophages after 48 h of conidial exposure. Notably, *A. fumigatus* strains elicited a graded, strain-specific cytokine profile, mirroring previous observations in *A. fischeri* [32]. When both species are considered jointly, there is considerable overlap in their cytokine signatures (Figure 5B).

We next examined the relationship between macrophage killing efficiency and cytokine levels. Among *A. fumigatus* strains, cytokine levels were positively correlated with macrophage killing efficiency (Figure S8): for instance, strains whose spores were highly resistant to macrophage killing (A1163 and JD8) provoked strong cytokine responses, whereas the melanin-deficient strain IP24 was susceptible and elicited a weak response [73]. Comparing these data with equivalent data from *A. fischeri* [32] showed that only two *A. fischeri* strains (CBS 150753 and CBS 150756) and one *A. fumigatus* strain (JD16) broke this trend, exhibiting substantially lower cytokine responses than their intermediate conidial survival would predict (Figure 5B). Importantly, both the cytokine levels themselves and their coupling to macrophage killing overlapped extensively between the two species. Thus, the inter-strain variance observed under *in vitro* coculture with murine macrophages blurs the boundaries of the two species, a finding that sharply contrasts with our genomic, phenotypic (from monoculture assays), and metabolic data, which consistently separated the species (Figs 1–4, S6).

### The virulence profiles of *A. fumigatus* and *A. fischeri* overlap in an immunocompromised murine model of pulmonary aspergillosis

To measure the pathogenic potential of each *A. fumigatus* strain *in vivo*, we next turned to a chemotherapeutic murine model of pulmonary aspergillosis (Methods). Using exactly the same methodology as for *A. fischeri* [31,32] enabled a direct comparison of the two species’ pathogenic potentials.

The lethalities of the reference strains of each species showed the expected strong demarcation between high and low lethality (Figure 5C, left and subpanel ***i***). Eleven *A. fumigatus* strains proved 100% lethal, including the melanin deficient strain (IP24) that appeared highly susceptible to macrophages. Among these fully lethal strains, we observed a wide range of mean time to death, from just under four days (DI19-230) to nine days (IP49) (Figure 5C, left) with several strains displaying disease progression rates greater than A1163 (Figure 5C, subpanel ***ii***). Strikingly, we found that the virulence profiles of three *A. fumigatus* strains overlapped with those of *A. fischeri* (Figure 5C, subpanel ***iii*** and ***iv***), with two strains (AFR9 and JD31) being among the least lethal strains of both species. Moreover, the most lethal *A. fischeri* strain (CBS 150757) appeared remarkably aggressive in the early stages of infection and produced greater mortality over the first seven days than did most *A. fumigatus* strains over the same period. These results are even more provocative when we consider that all but one *A. fumigatus* strain was a clinical isolate, whereas all *A. fischeri* strains were environmental isolates.

The overlap between *A. fumigatus* and *A. fischeri* in these *in vivo* results mirrored what we saw in macrophage cocultures, again revealing that traits can transcend species boundaries when strainwise variability is explored. Notably, per-strain lethality did not appear to be correlated to the *in vitro* metrics (Figure S9). There are examples of low-lethality strains that were highly susceptible to macrophages (JD31) and low-lethality strains that were only modestly susceptible to macrophages (AFR9 and JD48). Even some highly lethal strains were nevertheless very susceptible to killing my macrophages (DI19-230 and IP24). Consequently, strain heterogeneity can result in interspecific overlap along multiple axes (Figs 5D, S10, S11, S12, S13), suggesting that the early and late stages of fungal infection may be influenced by different aspects of the pathogen’s biology.

### Lack of association between gene content and virulence

We next sought to relate the *in vivo* results back to genomic features that have been previously suggested to correlate with virulence in *Aspergillus,* namely virulence-associated gene presence (Figure 3A), and accessory gene content (Figure S2). We first individually tested every gene present in at least one strain of each species (10,123 genes) for association with the mean host lifespan in our murine model of fungal disease. This resulted in 402 nominally significant genes (p < 0.05, Mann Whitney U test). The direction of effect of these associations was evenly split between increased- and decreased-lethality (208 vs 194 genes, respectively). Of those 402 genes, 197 could be related back to annotated *A. fumigatus* genes while the rest lacked functional annotation; only one of the 402 genes (AFU8G01222, *tmpL*) was part of the virulence-associated gene set. The TmpL protein has domain and structural similarity to several bacterial NRPSs, and its orthologs have been shown to be critical for oxidative stress homeostasis [74,75]. None of those associations remained significant following multiple testing corrections.

Accessory gene content has also been linked to virulence differences among strains of *A. fumigatus* [38]. We therefore tested for correlations between accessory gene content of the strains and lethality in mice. We tested each species separately, and both species together, and neither test suggested a significant relationship between the number of accessory genes and observed lifespan post infection (Spearman rho = -0.182, p = 0.516 and rho = -0.175, p= 0.175 in *A. fumigatus* and *A. fischeri* respectively). This lack of significant gene-trait associations supports a model of pathogenicity that transcends the boundaries of individual species.

## DISCUSSION

Comprehensive characterization of genetic, metabolomic, and phenotypic diversity of a diverse sample of strains from the pathogenic fungus *A. fumigatus* and comparison to diverse strains from its non-pathogenic close relative *A. fischeri* revealed that opportunistic pathogenicity can extend beyond the boundaries of individual species. While phenotypic overlap was apparent in several of the traits we measured (*e.g.,* Figure 1), it was most pronounced when the strains were challenged with either macrophages or in an *in vivo* murine model of pulmonary aspergillosis where at least two clinical strains of *A. fumigatus* exhibited lower virulence in mice than all but three *A. fischeri* strains (Figure 5). This result directly challenges the binary classification of fungi capable of opportunistically infecting humans as pathogenic or non-pathogenic, and suggests that pathogenic potential can sometimes exhibit greater variation within a species than between closely related species.

The observation that strain-level variance within each species can rival or even exceed interspecific variance during host-pathogen interactions raises the question of how specific genomic, metabolic, and phenotypic traits contribute to pathogenic potential. In our study, no single gene associations with virulence were statistically significant, nor were there any associations between virulence and pangenome accessory gene content. BGC and metabolite content similarly showed no clear relationship to degree of virulence, with similar profiles shared among the most- and least-lethal strains. Of the 207 virulence-associated genes surveyed, nearly all were present in both species with *A. fischeri* strains being reliably distinguished only by their shared lack of three genes in the fumagillin BGC. Even the *A. fumigatus* strain containing fewest virulence-associated genes (DI19_230 with 201 genes) was maximally lethal while the least lethal strains (AFR9, JD-31, and JD-16) contained 206 genes each. Therefore, the lack of clear, virulence-associated features among strains or between species suggests that pathogenicity is genetically complex, and possibly even heterogeneous, with different strains potentially achieving similar virulence outcomes through different routes (*i.e.,* through different combinations of traits).

### Geography, ecological opportunity, and pathogenic potential

Our results support a model where diversity among strains of species that are putatively either pathogenic or non-pathogenic can produce outcomes that defy species-level expectations of virulence. We propose that during fungal infections a given strain’s apparent virulence emerges from two sets of graded properties (Figure 6A): 1) the strain’s pre-existing amenability for growth in a mammalian host environment (ecological fitting; *e.g.,* thermotolerance to facilitate growth at mammalian body temperatures), and 2) the host’s ability to effectively forestall fungal infections (environmental filtering). This interaction between a given strain’s fit and a given host’s filtering determines the virulence of that strain in each host, with multiple strain-host combinations then occupying unique positions on a virulence landscape. Across this landscape, a strain’s virulence can range from benign colonization to lethal disease. Strains resulting in consistently high levels of virulence will be more frequently observed in clinical cases, and this rate of clinical ascertainment forms the basis for ascribing a degree of pathogenic potential to that strain. In the case of *A. fumigatus* and *A. fischeri*, while the former species is identified much more often in clinical settings, the full range of virulence of strains of both species suggests that the pathogenic potential of the latter may be underestimated. Importantly, because virulence is an emergent property of a complex host-pathogen interaction, individual fungal genes or traits are not expected to be strongly predictive of clinical outcome.

**Figure 6.**
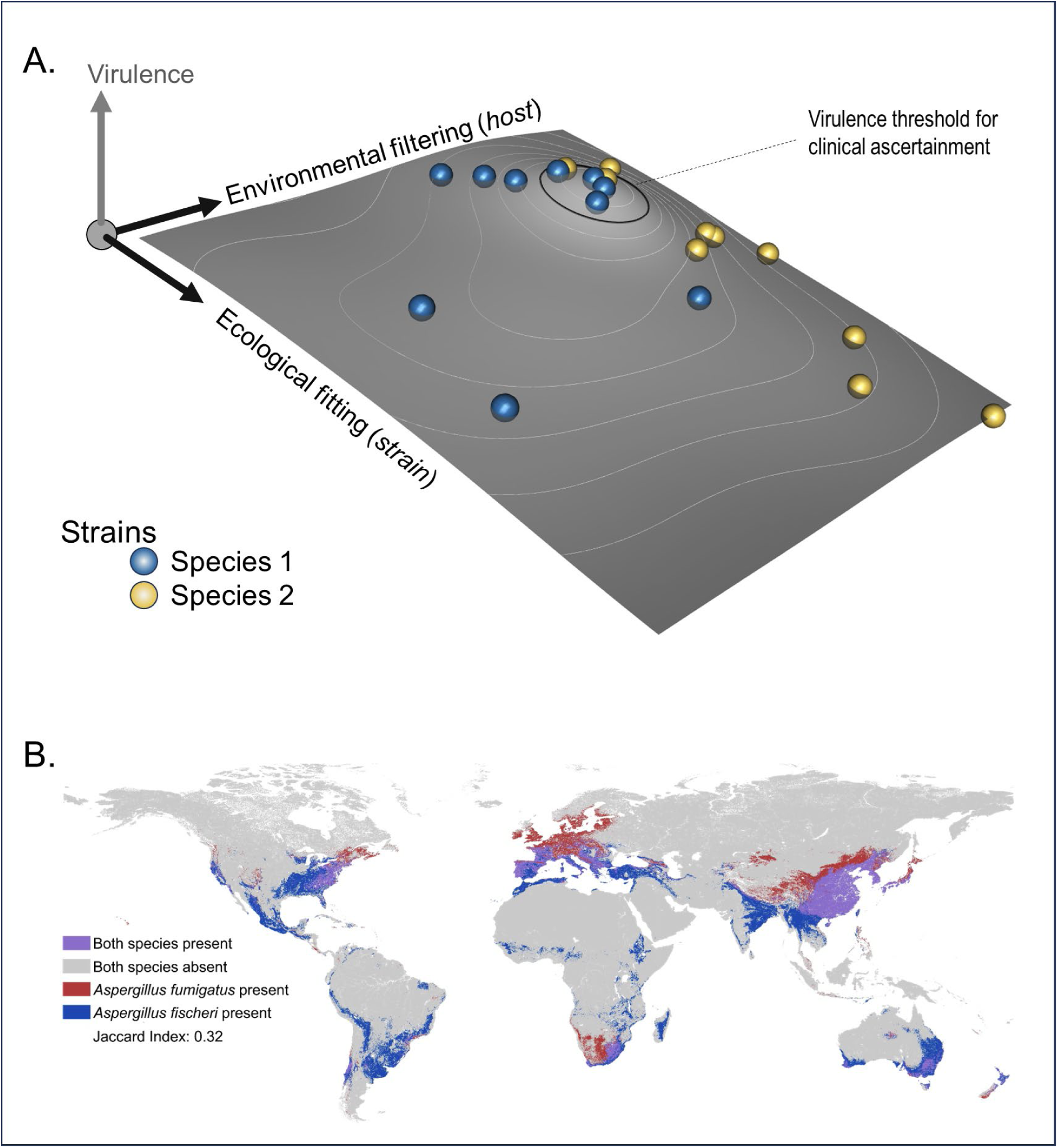
Fungal pathogenicity as an ecological process. **A.** The degree of virulence of any given strain (ball) is an emergent property of the interaction of two ecological processes: (1) **ecological fitting**, where naturally evolved traits of a given fungal strain coincidentally permit growth in a host, and (2) **environmental filtering**, where a given host’s immune responses select or filter for which strains can persist. Considered across all strains and hosts, the virulence landscape (surface) of a fungal species (yellow or blue) is determined by each strain’s ecological fitting and the permissiveness of the host environment. The clinical classification of any species as “pathogenic” is determined by how frequently strains of the species exceed this threshold. **B.** Predicted global distribution of two *Aspergillus* species. Machine learning model predictions of species ranges for *A. fumigatus* and *A. fischeri* provide some context for the **ecological opportunity** of these species (data from [76]). An important limitation of these predictions is that they rely strongly on broad-scale environmental metrics and do not account for susceptible hosts or any artificial microenvironments that a clinical setting may provide.

Implicit in this formulation of pathogenic potential is the notion that the diversity and constitution of co-occurring host and pathogen populations will influence the classification of any species as “pathogenic”. In other words, ecological opportunity is a prerequisite for any position on the host impact landscape to be realized. For example, our predictive model for species distribution [76](Figure 6B) reveals that both *A. fumigatus* and *A. fischeri* have broader global ranges than previously appreciated, yet show only limited areas of large-scale geographic overlap. This implies that the ecological opportunity for host encounter is neither uniform nor equivalent between species, and may vary substantially by region. Moreover, because clinical identification of *Aspergillus species* typically relies only on morphological or ITS-based methods, some cases of aspergillosis currently attributed to *A. fumigatus* may in fact involve cryptic species, including *A. fischeri* [77]. We hypothesize that more rigorous molecular identification of clinical isolates would reveal such misattribution, particularly in regions where only *A. fischeri* is present or in regions where both species co-occur.

## CONCLUSION

Our findings carry several implications for how the evolution of opportunistic fungal pathogenesis can be better explored going forward. First, monoculture phenotyping of fungi must continue to be broadened to encompass a wider diversity of conditions and strains rather than relying on assumptions derived from one or two reference genotypes. Fungal traits that contribute to opportunistic pathogenicity evolve in diverse natural environments and only coincidentally facilitate host colonization; understanding how these traits co-vary across strains is essential to understanding how different lineages exhibit different pathogenic potentials. Similarly, species surveillance efforts must become geographically broader and taxonomically more sensitive, particularly when surveying clinically associated niches. Clinical cases can be misattributed to the most common species due to insufficient diagnostic protocols; this scenario is probably more common than is currently appreciated and likely extends to several cryptic *Aspergillus* species. A deeper understanding of the ecological niches occupied by representative strains of each species is essential for better predicting which lineages are most likely to opportunistically infect humans.

Second, to become a successful pathogen, an ecologically fit fungus must ultimately survive environmental filtering by the host. This is a dynamic process involving both early and late immune responses, in host environments that are themselves heterogeneous. A more complete understanding of the genetic and immunologic factors governing antifungal immunity is needed to capture the dynamics of this second component of virulence. One such approach could involve large-scale integration of electronic health records that include host genotype data, enabling better characterization of how host variation shapes infection outcomes. Ultimately, pathogenic potential is a latent property of the host–pathogen interaction rather than a fixed attribute of any single species or strain; thus, moving beyond species-level assumptions toward detailed characterization of individual clinically relevant fungal isolates will be necessary to anticipate and mitigate the growing threat of invasive fungal disease.

## METHODS

### *A. fumigatus* strains analyzed in this study

All strains of *A. fumigatus* analyzed here were provided by the laboratory of Dr. John Gibbons (University of Massachusetts, Amherst). Strain identifiers and metadata are provided in Table 1. Population analysis of *A. fumigatus* strains

Whole-genome Illumina sequence data for all isolates were adapter- and quality-filtered using Trim Galore, mapped to the *A. fumigatus* Af293 reference genome with BWA-MEM, and SNPs were called using GATK HaplotypeCaller following the germline short variant discovery best practices pipeline[78–80]. A final set of 7,386 informative SNPs was obtained from 214,959 raw SNPs after filtering with VCFtools for minor allele frequency (MAF < 5%), missing genotype rate (>20%), and minimum inter-SNP spacing (3,500 bp) to reduce the effects of physical linkage. A maximum-likelihood phylogenetic tree was constructed from the 7,386-SNP alignment using FastTree[78] v2.1.10 with a generalized time-reversible (GTR) substitution model, with local support values assessed via the Shimodaira-Hasegawa test, and the tree was visualized using the R package ggtree. Population structure was inferred using ADMIXTURE[81] v1.3 across K = 1–85, where K represents the number of populations, with the optimal K determined by 5-fold cross-validation as the value minimizing cross-validation error.

### Genome sequencing and quality assessment

Each *A. fumigatus* strain was grown at 30°C prior to DNA extraction. For long read sequencing, high molecular weight genomic DNA was isolated from pellets and sequenced by Oxford Nanopore (ONT) by SeqCenter (Pittsburgh, PA, USA). Short read DNA sequences were downloaded from NCBI SRA.

### De *novo* genome assembly

ONT reads from each strain were checked for contamination using Kaiju (v1.8.2) [82] and assembled using Flye (v.2.9-b1774) [83]. Long-read assemblies were polished with the Illumina reads using *pilon* (v1.24) [84]. The resulting assemblies were evaluated for completeness using BUSCO (fungi_odb10 and eurotiomycetes_obd10) [85]. Assembly parameters were optimized for each strain so as to maximize BUSCO completeness. Optimal assemblies were obtained by thresholding Flye for a read error of 0.02, followed by 2 rounds of Pilon polishing. Assemblies were evaluated for quality using QUAST (QUality ASsessment Tool; v.5.2.0) [86] and the best assembly (DTO15) was chosen to serve as a reference genome for all the other strains.

### Phylogenetics

To infer the strain phylogeny, single copy orthologs were extracted from 16 *A. fumigatus*, 16 *A. fischeri* , and two outgroup genomes (*Aspergillus lentulus*, DTO_433_A3_HYB and *Aspergillus fumisynnematus*, DTO_171-B2_HYB) using BUSCO (v. 5.4.6) [85] and the eurotiomycetes_odb10 data base (N=3,546 BUSCO entries). After filtering for BUSCO genes present across at least 95% of the samples and outgroups (N=3135) each gene was individually aligned with MAFFT. The resulting alignments were concatenated and used to build the species and strain phylogeny with IQ-TREE (v. 2.2.2.3) [87].

### Pangenome analysis

The *A. fumigatus* pangenome was constructed using Pantools (v4.1.1) [88,89]. In short, a database was created using the amino acid sequences for each of the gene models produced by Funnanotate for each of the *A. fumigatus* strains. Initial orthogroups were determined using the Pantools’ build_panproteome function, then the optimal_grouping function identified the optimal sequence similarity threshold (75%) for maximizing precision and recall (the F-score). Initial assessment of core, accessory, and singleton status of orthogroups was performed using the gene_classification function with default settings. The gene_classification function also provided a Heap’s Law calculation.

Next, orthogroups from Pantools were clustered by sequence similarity with a dynamic similarity threshold using ALFATClust (version available as of 04-10-2023) with default settings (Chiu & Ong 2022). This resulted in a set of 14754 gene clusters across all 16 strains. Each of these gene clusters were assigned a number within the parent orthogroup. Gene clusters were characterized as core, accessory, or singletons based on the number of strains represented in each cluster. Core genes were those gene clusters that contained sequences from all 16 strains, accessory genes contained sequences from 2-15 strains, and genes found in a single strain were considered singletons. Gene copy number was assessed by counting the number of sequences from each strain within a gene cluster. In downstream analysis, multicopy gene clusters (gene clusters that contained multiple genes from a single strain) and gene clusters whose average percent identity was lower than 70% were excluded from analysis.

To relate *A. fumigatus* pangenes back to the canonical reference strains of *A. fumigatus* (Af293) and *A. fischeri* (NRRL 181), we sourced annotations for Af293 and NRRL 181 from FungiDB release 62 [90]. These reference sequences were used to construct a custom BLAST database against which, pangenecluster centroids (identified using DIAMOND (v2.1.6) [91]) were reciprocally BLASTed Any duplicate BLAST results were resolved by selecting for maximum bit score. The homologous genes found were used for annotating the pangenome.

### Biosynthetic Gene Cluster detection

To identify biosynthetic gene clusters (BGCs), we collected the genes and locus tags from primary literature and the BGC repository MIBiG (v3.1) [92]. BGCs in the *A. fumigatus* strains were annotated by antiSMASH (v6.0.1) [93] and manually reviewed to confirm gene content, borders and syntenic similarity.

### Growth and extraction of *Aspergillus fumigatus* cultures for analyses of secondary metabolite profiles

*A. fumigatus* strains (n=16) were grown on oatmeal media (Quaker breakfast oatmeal) for chemical characterization, since previous studies showed that *Aspergillus* sp. biosynthesized a larger set of secondary metabolites on this media[33]. Briefly, fungal cultures were started by aseptically cutting mycelia from a culture growing in a Petri dish on malt extract agar (MEA, Difco), and this material was transferred to a sterile Falcon tube with 10 mL of liquid YESD media (YESD; 20 g soy peptone, 20 g dextrose, 5 g yeast extract, 1 L distilled H_2_O). This seed culture was grown for 7 days on an orbital shaker (∼100 rpm) at room temperature (∼23°C) and used to inoculate the oatmeal cereal media, which was prepared by adding 10 g of oatmeal in a 250 mL, wide mouth Erlenmeyer flask with ∼17 mL of deionized water followed by autoclaving at 121°C for 30 min. For each of the 16 strains, three flasks of each of the samples (i.e., three biological replicates per sample per growth temperature) were incubated for 10 d at both 30°C and 37°C. Since it is known that light modulates growth, sexual reproduction, and secondary metabolite production in fungi[94], the incubators were equipped with LED lamps (Ustellar, flexible LED strip lights; 24 W) with light regulated into 12:12 h dark/light cycles[95].

At the end of 10 d, each individual flask was extracted by adding 70 mL of CHCl_3_-MeOH (1:1), chopped with a spatula, and shaken overnight (∼16 h) at ∼100 rpm at room temperature. The cultures were then filtered *in vacuo*, and 90 mL of CHCl_3_ and 150 mL of deionized H_2_O were added to each of the filtrates. The mixture was then transferred to a separatory funnel and shaken vigorously. The organic layer (*i.e*., bottom layer) was drawn off and evaporated to dryness *in vacuo*. This dried material was reconstituted in 100 mL of CH_3_CN-MeOH (1:1) and 100 mL of hexanes, transferred to a separatory funnel, and shaken vigorously. The defatted organic layer was evaporated to dryness.

All the defatted organic layers from each biological replicate at both 30°C and 37°Cwere analyzed individually by UPLC-HRESI-MS utilizing a Thermo LTQ Orbitrap XL mass spectrometer equipped with an electrospray ionization source. A Waters Acquity UPLC was utilized with a BEH C_18_ column (1.7 μm; 50 mm × 2.1 mm) set to 40°C and a flow rate of 0.3 mL/min. The mobile phase consisted of a linear gradient of CH_3_CN- H_2_O (both acidified with 0.1% formic acid), starting at 15% CH_3_CN and increasing linearly to 100% CH_3_CN over 8 min, with a 1.5 min hold before returning to the starting conditions. The profile of secondary metabolites of each biological replicate was examined using well-established procedures for metabolite identification[59,60], similar to previous studies [33,96].

### Principal component analysis of metabolomics results

Principal component analysis (PCA) and hierarchical clustering were performed on the UPLC-HRESI-MS data. Untargeted UPLC-HRESI-MS datasets for each sample were individually aligned, filtered, and analyzed using MZmine (v2.53) (https://sourceforge.net/projects/mzmine/). Peak list filtering and retention time alignment algorithms were used to refine peak detection, and the join algorithm integrated all sample profiles into a data matrix using the following parameters: Mass detection: MS1 positive mode. ADAP: Group intensity threshold: 20000, Min highest intensity: 60000, m/z tolerance: 0.003.

Chromatogram deconvolution: Wavelets (ADAP) algorithm. Join aligner and gap filling: retention time tolerance: 0.05 min, m/z tolerance: 0.0015 m/z. The resulting data matrix was exported to Excel (Microsoft) for analysis as a set of m/z–RT (retention time) pairs with individual peak areas. Samples that did not possess detectable quantities of a given marker ion were assigned a peak area of zero to maintain the same number of variables for all sample sets. Ions that did not elute between 1 and 10 min and/or had an m/z ratio < 200 or > 900 Da were removed from analysis. Relative standard deviation was used to understand the quantity of variance between the injections, which may differ slightly based on instrument variance. A cutoff of 1.0 was used at any given m/z–RT pair across the biological replicate injections, and if the variance was greater than the cutoff, it was assigned a peak area of zero. PCA analysis and hierarchical clustering were created with Python. The PCA scores plots were generated using the averaged data of the three individual biological replicates.

### Morgan fingerprint generation

Morgan fingerprints were generated for each compound from its molecular structure file (.mol) using the RDKit cheminformatics toolkit (RDKit: Open-source cheminformatics. https://www.rdkit.org). Each detected molecule was loaded via the `Chem.MolFromMolFile()` function, and a count-based Morgan fingerprint (count vector) was computed using the `GetMorganGenerator` method with a radius of 2 and a fingerprint length of 2048 bits. The `GetCountFingerprint()` function was used to generate the fingerprint as a vector of integer counts for each bit position, rather than a binary presence/absence vector. All fingerprint generation was performed in Python (version 3.11). For summarizing all molecules detected in a strain (*e.g.,* Figure 4A) individual fingerprints were summed across all positions.

### Murine model of pulmonary aspergillosis

Wild-type BALB/c female mice, body weight 20 to 22 g, aged 8-9 weeks, were kept in the Animal Facility of the Laboratory of Molecular Biology of the School of Pharmaceutical Sciences of Ribeirão Preto, University of São Paulo (FCFRP/USP), in a clean and silent environment, under normal conditions of humidity and temperature, and with a 12:12 h dark/light cycle. The mice were given food and water *ad libitum* throughout the experiments. Mice were immunosuppressed with cyclophosphamide (150 mg per kg of body weight), which was administered intraperitoneally on days -4 and -1 (pre-infection) and +2 days (post infection). Hydrocortisone Acetate (200mg/ kg body weight) was injected subcutaneously on day -3. *A. fumigatus* strains and the *A. fumigatus* A1163 strain (a clinically derived and highly virulent strain, that we used as a positive control), were grown on minimal media for 2 days prior to infection. Fresh conidia were harvested in PBS and filtered through a Miracloth (Calbiochem). Conidial suspensions were spun for 5 min at 3,000 x g, washed three times with PBS, counted using a hemocytometer, and resuspended at a concentration of 5.0 x 10^6^ conidia/ ml. The viability of the administered inoculum was determined by incubating a serial dilution of the conidia on minimal media at 37°C. Mice were anesthetized by halothane inhalation and infected by intranasal instillation of 1.0 x 10^5^ conidia in 20 µl of PBS. As a negative control, a group of 10 mice received PBS only. Mice were weighed every 24 h from the day of infection and visually inspected twice daily. The statistical significance of comparative survival values was calculated using the Log-rank (Mantel-Cox) Test and Gehan-Breslow-Wilcoxon tests, as implemented in the Prism statistical analysis package.

The virulence of each *A. fumigatus* strain was evaluated in groups of 10 infected mice. The reference strain *A. fumigatus* A1163 was included as a positive control group (10 infected mice). A single group of 10 PBS-treated mice served as the negative control for all experimental groups within each replicate. All experiments were performed in two independent biological replicates.

The principles that guide our studies are based on the Declaration of Animal Rights ratified by UNESCO on January 27, 1978 in its 8th and 14th articles. All protocols adopted in this study were approved by the local ethics committee for animal experiments from the University of São Paulo, Campus of Ribeirão Preto (Permit Number: 08.1.1277.53.6; Studies on the interaction of *A. fumigatus* with animals). Groups of five animals were housed in individually ventilated cages and were cared for in strict accordance with the principles outlined by the Brazilian College of Animal Experimentation (COBEA) and Guiding Principles for Research Involving Animals and Human Beings, American Physiological Society. All efforts were made to minimize suffering. Animals were clinically monitored at least twice daily and humanely sacrificed if moribund (defined by lethargy, dyspnea, hypothermia, and weight loss). All stressed animals were sacrificed by cervical dislocation.

### Macrophage cytokine response to conidia

*Aspergillus* strains were cultivated on minimal medium agar plates at 37°C for 3 days. Conidia were harvested in sterile water with 0.05 % (vol/vol) Tween20. The resulting suspension was filtered through two layers of gauze (Miracloth, Calbiochem). The conidial concentration was determined using a hemocytometer.

BALB/c bone marrow-derived macrophages (BMDMs) were obtained as previously described [97]. Briefly, bone marrow cells were cultured for 7–9 days in RPMI 20/30, which consists of RPMI-1640 medium (Gibco, Thermo Fisher Scientific Inc.), supplemented with 20 % (vol/ vol) FBS and 30 % (vol/vol) L-Cell Conditioned Media (LCCM) as a source of macrophage colony-stimulating factor (M-CSF) on non-treated Petri dishes (Optilux - Costar, Corning Inc. Corning, NY). Twenty-four hours before experiments, BMDM monolayers were detached using cold phosphate-buffered saline (PBS) (Hyclone, GE Healthcare Inc. South Logan, UT) and cultured, as specified, in RPMI-1640 (Gibco, Thermo Fisher Scientific Inc.) supplemented with 10 % (vol/vol) FBS, 10 U/mL penicillin, and 10 µg/mL streptomycin, (2 mM) L-glutamine, (25 mM) HEPES, pH 7.2 (Gibco, Thermo Fisher Scientific Inc.) at 37°C in 5 % (vol/vol) CO_2_ for the indicated periods.

BMDMs were seeded at a density of 10^6^ cells/ml in 24-well plates (Greiner Bio-One, Kremsmünster, Austria). The cells were challenged with the conidia of different strains at a multiplicity of infection of 1:10 and incubated at 37°C with 5 % (vol/vol) CO_2_ for 48h. BMDMs were also stimulated with lipopolysaccharide (LPS; standard LPS, *E. coli* O111:B4; Sigma-Aldrich, 500 ng/mL) plus Nigericin (tlrl-nig, InvivoGen 5 μg/mL) and cell medium alone, which were used respectively as the positive and negative controls. Cell culture supernatants were collected and stored at −80°C until they were assayed for TNF-α, IL-1β, and IL-6 release using Mouse DuoSet ELISA kits (R&D Systems, Minneapolis, MN, USA, according to the manufacturer’s instructions. For cytokine determination, plates were analyzed by using a microplate reader (Synergy™ HTX Multi-Mode, BioTek) measuring absorbance at 450 nm. Cytokine concentrations were interpolated from a standard curve and statistical significance was determined using an ANOVA.

### Macrophage killing assay

BMDMs were seeded at a density of 10^6^ cells/ml in 24-well plates (Corning^®^ Costar^®^) and were challenged with conidia at a multiplicity of infection of 1:10 and incubated a 37°C with 5 % (vol/vol) CO_2_ for 24h. After incubation media was removed the cells were washed with ice-cold PBS and finally 2 ml of sterile water was added to the wells. A P1000 tip was then used to scrape away the cell monolayer and the cell suspension was collected. This suspension was then diluted 1:1000 and 100 μl was plated on Sabouraud agar before the plates were incubated at 37°C overnight and the colonies were counted. 50 μl of the inoculum adjusted to 10^3^/ml was also plated on SAB agar to correct CFU counts. The CFU/ml for each sample was calculated and compared to the A1163 wild-type strain using (GraphPad Prism 8.0, La Jolla, CA). All assays were performed in four replicates in two independent experiments.

### Growth phenotyping

#### Strains, media, and phenotypic assays

*A. fumigatus* and *A. fischeri* strains (Table 1) were grown in solid minimal medium (MM; 1% [wt/vol] glucose, 50 mL of a 20x salt solution, 0.1% [vol/vol] trace elements, 2% [wt/vol] agar, pH 6.5) at 37°C or a rich medium (YAG; 2% [wt/vol] glucose, 0.5% [wt/vol] yeast extract, 0.1% [vol/vol] traces elements; 2% [wt/vol] agar). The trace elements and nitrate salts were prepared as previously described (Kafer, 1977). For the liquid medium, the agar was not added. For phenotypic characterization, plates containing solid MM or YAG were centrally inoculated with 5 µL of a 1 × 10^6 conidia suspension of each strain and incubated at 30, 37, or 44°C. Additionally, plates containing solid MM were centrally inoculated with 5 µL of a 1 × 10^6 conidia suspension of each strain in the presence or absence of various stressor agents (Congo red 50 µg/mL; calcofluor white- CFW 25 µg/mL; menadione 0.02mM; H_2_O_2_ 3mM; T-tubyl 0.85mM) and incubated at 37°C. After 120 h of incubation, radial growth was measured. All radial growth analyses were performed in triplicate; mean ± standard deviation (SD) is reported.

#### Staining and labeling of cell surface components

Cell wall surface polysaccharide staining was performed as described previously [98]. Briefly, 10^4^ conidia (swollen and resting conidia) for *A. fumigatus* strains were inoculated in 200 µL of liquid MM and incubated for 4 h at 37°C (swollen conidia) and 4°C (resting conidia) before the culture medium was removed and conidia were UV-irradiated (600,000 mJ). For dectin labeling, 200 µL of a blocking solution (2% [vol/vol] goat serum, 1% [vol/vol] bovine serum albumin [BSA], 0.1% [vol/vol] Triton X-100, 0.05% [vol/vol] Tween 20, 0.05% [vol/vol] sodium azide, and 0.01 M PBS) was added to each well. Samples were incubated for 30 min at room temperature (RT), and 0.2 µg/mL of Fc-h-dectin-hFc (Invivogen) was added to the UV-irradiated conidia and incubated for 1 h at RT, followed by the addition of 1:1,000 DyLight 594-conjugated goat anti-human IgG1 (Abcam) for 1 h at RT. Conidia were then washed with phosphate-buffered saline (PBS), and fluorescence was read at 587 nm excitation and 615 nm emission. For chitin staining, 200 µL of a PBS solution containing 10 mg/mL calcofluor white (CFW) was added to the UV-irradiated conidia, which were incubated for 5 min at RT and washed with PBS before fluorescence was measured at 380 nm excitation and 450 nm emission. All experiments were performed with at least 4 replicates, and fluorescence was measured using a microtiter plate reader (Synergy HTX Multi-Mode, BioTek).

#### Fungal adhesion and biofilm formation assays

To determine the adhesion capacity of conidia from the deleted mutant and A1163 strains, 1 × 10^4^ conidia were inoculated into 200 ul liquid MM in a 96-well polystyrene microtiter plate. Following an initial adherence phase of 4 h during static incubation at 37°C (swollen conidia) or 4°C (resting conidia), unbound conidia were washed with sterile PBS. Fresh liquid MM was added to the adhered conidia, and static submerged cultures were grown for up to 24 h at 37°C. Subsequently, the plate was washed exhaustively with PBS prior to incubation with 200 µl of 0.5% (wt/vol) crystal violet solution for 5 min at room temperature. The stained mycelia were then thoroughly washed with sterile water and air-dried. Finally, the crystal violet was eluted from the wells using 100% ethanol, and the absorbance was measured at 590 nm using a Synergy HTX Multimode Reader (Agilent Biotek).

## Data availability statement

Sequencing data and genome assemblies associated with this project will become available through GenBank upon publication. All other data necessary to replicate this work will be available on FigShare.

## Acknowledgements

We thank members of the Rokas lab for helpful discussions and feedback. Computational infrastructure at Vanderbilt University was provided by The Advanced Computing Center for Research and Education (ACCRE). This research was supported by the National Institutes of Health National Institute of Allergy and Infectious Diseases (R01 AI153356). O.L.R. was supported by the National Science Foundation Graduate Research Fellowship Program under Grant No. 2444112. Research in A.R.’s lab is also supported by the National Science Foundation (DEB-2110404) and Vanderbilt University. The funders had no role in study design, data collection, and analysis, decision to publish, or preparation of the manuscript.

## REFERENCES

1. Bongomin F, Gago S, Oladele RO, Denning DW. Global and Multi-National Prevalence of Fungal Diseases-Estimate Precision. J Fungi (Basel). 2017;3. doi:10.3390/jof3040057

2. Hibbett D, Nagy LG, Nilsson RH. Fungal diversity, evolution, and classification. Curr Biol. 2025;35: R463–R469.

3. Hawksworth DL, Lücking R. Fungal Diversity Revisited: 2.2 to 3.8 Million Species. Microbiol Spectr. 2017;5. doi:10.1128/microbiolspec.FUNK-0052-2016

4. Sherrington SL, Kumwenda P, Kousser C, Hall RA. Host sensing by pathogenic fungi. Adv Appl Microbiol. 2018;102: 159–221.

5. Rokas A. Evolution of the human pathogenic lifestyle in fungi. Nature Microbiology. 2022;7: 607–619.

6. Janzen DH. On Ecological Fitting. Oikos. 1985;45: 308.

7. Janzen DH. When is it coevolution? Evolution. 1980;34: 611–612.

8. Agosta SJ, Klemens JA. Ecological fitting by phenotypically flexible genotypes: implications for species associations, community assembly and evolution: Ecological fitting. Ecol Lett. 2008;11: 1123–1134.

9. Kraft NJB, Adler PB, Godoy Ó, James EC, Fuller S, Levine JM, et al. Community assembly, coexistence and the environmental filtering metaphor. Funct Ecol. 2015;29: 592–599.

10. Robert VA, Casadevall A. Vertebrate endothermy restricts most fungi as potential pathogens. J Infect Dis. 2009;200: 1623–1626.

11. André AC, Laborde M, Marteyn BS. The battle for oxygen during bacterial and fungal infections. Trends Microbiol. 2022;30: 643–653.

12. Puerner C, Vellanki S, Strauch JL, Cramer RA. Recent advances in understanding the human fungal pathogen hypoxia response in disease progression. Annu Rev Microbiol. 2023;77: 403–425.

13. Yaakoub H, Mina S, Calenda A, Bouchara J-P, Papon N. Oxidative stress response pathways in fungi. Cell Mol Life Sci. 2022;79: 333.

14. Chadwick BJ, Lin X. Effects of CO2 in fungi. Curr Opin Microbiol. 2024;79: 102488.

15. Vylkova S. Environmental pH modulation by pathogenic fungi as a strategy to conquer the host. PLoS Pathog. 2017;13: e1006149.

16. Weerasinghe H, Stölting H, Rose AJ, Traven A. Metabolic homeostasis in fungal infections from the perspective of pathogens, immune cells, and whole-body systems. Microbiol Mol Biol Rev. 2024;88: e0017122.

17. Cordero RJB, Casadevall A. Melanin. Curr Biol. 2020;30: R142–R143.

18. Lionakis MS, Drummond RA, Hohl TM. Immune responses to human fungal pathogens and therapeutic prospects. Nat Rev Immunol. 2023;23: 433–452.

19. Bhatt V, Saleem A. Review: Drug-induced neutropenia--pathophysiology, clinical features, and management. Ann Clin Lab Sci. 2004;34: 131–137.

20. Pana Z-D, Farmaki E, Roilides E. Host genetics and opportunistic fungal infections. Clin Microbiol Infect. 2014;20: 1254–1264.

21. Antinori S. New insights into HIV/AIDS-associated cryptococcosis. ISRN AIDS. 2013;2013: 471363.

22. Wang Y, Yu Y, Liu J, Li L, Chen X, Xiong S, et al. Intrinsic and extrinsic factors affecting the evolution of virulence in the HIV-associated opportunistic human fungal pathogen Cryptococcus neoformans. Virulence. 2025;16: 2546067.

23. Casadevall A. Cards of virulence and the global virulome for humans. Microbe Wash DC. 2006;1: 359–364.

24. Casadevall A, Fang FC, Pirofski L-A. Microbial virulence as an emergent property: consequences and opportunities. PLoS Pathog. 2011;7: e1002136.

25. Casadevall A, Pirofski L-A. The damage-response framework of microbial pathogenesis. Nat Rev Microbiol. 2003;1: 17–24.

26. Denning DW. Global incidence and mortality of severe fungal disease. Lancet Infect Dis. 2024. doi:10.1016/S1473-3099(23)00692-8

27. Steinbach WJ, Marr KA, Anaissie EJ, Azie N, Quan S-P, Meier-Kriesche H-U, et al. Clinical epidemiology of 960 patients with invasive aspergillosis from the PATH Alliance registry. J Infect. 2012;65: 453–464.

28. WHO fungal priority pathogens list to guide research, development and public health action. World Health Organization; 25 Oct 2022 [cited 27 May 2026]. Available: https://www.who.int/publications/i/item/9789240060241

29. Rokas A, Mead ME, Steenwyk JL, Oberlies NH, Goldman GH. Evolving moldy murderers: Aspergillus section Fumigati as a model for studying the repeated evolution of fungal pathogenicity. PLoS Pathog. 2020;16: e1008315.

30. Lamoth F. Aspergillus fumigatus-Related Species in Clinical Practice. Front Microbiol. 2016;7: 683.

31. Steenwyk JL, Mead ME, Knowles SL, Raja HA, Roberts CD, Bader O, et al. Variation Among Biosynthetic Gene Clusters, Secondary Metabolite Profiles, and Cards of Virulence Across Aspergillus Species. Genetics. 2020;216: 481–497.

32. Rinker DC, Sauters TJC, Steffen K, Gumilang A, Raja HA, Rangel-Grimaldo M, et al. Strain heterogeneity in a non-pathogenic Aspergillus fungus highlights factors associated with virulence. Commun Biol. 2024;7: 1082.

33. Mead ME, Knowles SL, Raja HA, Beattie SR, Kowalski CH, Steenwyk JL, et al. Characterizing the Pathogenic, Genomic, and Chemical Traits of Aspergillus fischeri, a Close Relative of the Major Human Fungal Pathogen Aspergillus fumigatus. mSphere. 2019;4. doi:10.1128/mSphere.00018-19

34. Mead ME, Steenwyk JL, Silva LP, de Castro PA, Saeed N, Hillmann F, et al. An evolutionary genomic approach reveals both conserved and species-specific genetic elements related to human disease in closely related Aspergillus fungi. Genetics. 2021;218. doi:10.1093/genetics/iyab066

35. Brown A, Mead ME, Steenwyk JL, Goldman GH, Rokas A. Extensive non-coding sequence divergence between the major human pathogen Aspergillus fumigatus and its relatives. Front Fungal Biol. 2022;3: 802494.

36. Gluck-Thaler E, Ralston T, Konkel Z, Ocampos CG, Ganeshan VD, Dorrance AE, et al. Giant Starship Elements Mobilize Accessory Genes in Fungal Genomes. Mol Biol Evol. 2022;39. doi:10.1093/molbev/msac109

37. Kowalski CH, Beattie SR, Fuller KK, McGurk EA, Tang Y-W, Hohl TM, et al. Heterogeneity among Isolates Reveals that Fitness in Low Oxygen Correlates with Aspergillus fumigatus Virulence. MBio. 2016;7. doi:10.1128/mBio.01515-16

38. Barber AE, Sae-Ong T, Kang K, Seelbinder B, Li J, Walther G, et al. Aspergillus fumigatus pan-genome analysis identifies genetic variants associated with human infection. Nat Microbiol. 2021;6: 1526–1536.

39. Gluck-Thaler E, Forsythe A, Puerner C, Gutierrez-Perez C, Stajich JE, Croll D, et al. Giant transposons promote strain heterogeneity in a major fungal pathogen. MBio. 2025;16: e0109225.

40. Lofgren LA, Ross BS, Cramer RA, Stajich JE. The pan-genome of Aspergillus fumigatus provides a high-resolution view of its population structure revealing high levels of lineage-specific diversity driven by recombination | PLOS Biology. PLoS Biol. 2022;20: e3001890.

41. Horta MAC, Steenwyk JL, Mead ME, Dos Santos LHB, Zhao S, Gibbons JG, et al. Examination of genome-wide ortholog variation in clinical and environmental isolates of the fungal pathogen Aspergillus fumigatus. MBio. 2022;13: e0151922.

42. Estabrook GF, McMorris FR, Meacham CA. Comparison of undirected phylogenetic trees based on subtrees of four evolutionary units. Syst Zool. 1985;34: 193.

43. Samson RA, Hong S, Peterson SW, Frisvad JC, Varga J. Polyphasic taxonomy of Aspergillus section Fumigati and its teleomorph Neosartorya. Stud Mycol. 2007;59: 147–203.

44. Sugui JA, Kwon-Chung KJ, Juvvadi PR, Latgé J-P, Steinbach WJ. Aspergillus fumigatus and related species. Cold Spring Harb Perspect Med. 2014;5: a019786.

45. Arico-Muendel C, Centrella PA, Contonio BD, Morgan BA, O’Donovan G, Paradise CL, et al. Antiparasitic activities of novel, orally available fumagillin analogs. Bioorg Med Chem Lett. 2009;19: 5128–5131.

46. Lind AL, Wisecaver JH, Lameiras C, Wiemann P, Palmer JM, Keller NP, et al. Drivers of genetic diversity in secondary metabolic gene clusters within a fungal species. PLoS Biol. 2017;15: e2003583.

47. Fallon JP, Reeves EP, Kavanagh K. Inhibition of neutrophil function following exposure to the Aspergillus fumigatus toxin fumagillin. J Med Microbiol. 2010;59: 625–633.

48. Guruceaga X, Ezpeleta G, Mayayo E, Sueiro-Olivares M, Abad-Diaz-De-Cerio A, Aguirre Urízar JM, et al. A possible role for fumagillin in cellular damage during host infection by Aspergillus fumigatus. Virulence. 2018;9: 1548–1561.

49. Lowther WT, Matthews BW. Structure and function of the methionine aminopeptidases. Biochim Biophys Acta. 2000;1477: 157–167.

50. O’Hanlon KA, Cairns T, Stack D, Schrettl M, Bignell EM, Kavanagh K, et al. Targeted disruption of nonribosomal peptide synthetase pes3 augments the virulence of Aspergillus fumigatus. Infect Immun. 2011;79: 3978–3992.

51. Eagan JL, Keller NP. Fungal secondary metabolism. Curr Biol. 2025;35: R503–R508.

52. Raffa N, Keller NP. A call to arms: Mustering secondary metabolites for success and survival of an opportunistic pathogen. PLoS Pathog. 2019;15: e1007606.

53. Latif H, Gross M, Fischer D, Lierz M, Usleber E. Immunochemical analysis of fumigaclavine mycotoxins in respiratory tissues and in blood serum of birds with confirmed aspergillosis. Mycotoxin Research. 2015;31: 177–183.

54. Wadhwa K, Kapoor N, Kaur H, Abu-Seer EA, Tariq M, Siddiqui S, et al. A Comprehensive Review of the Diversity of Fungal Secondary Metabolites and Their Emerging Applications in Healthcare and Environment. Mycobiology. 2024; 335–387.

55. Patel R, Hossain MA, German N, Al-Ahmad AJ. Gliotoxin penetrates and impairs the integrity of the human blood-brain barrier in vitro. Mycotoxin Res. 2018;34: 257–268.

56. Lewis RE, Wiederhold NP, Chi J, Han XY, Komanduri KV, Kontoyiannis DP, et al. Detection of gliotoxin in experimental and human aspergillosis. Infect Immun. 2005;73: 635–637.

57. Rokas A, Wisecaver JH, Lind AL. The birth, evolution and death of metabolic gene clusters in fungi. Nat Rev Microbiol. 2018;16: 731–744.

58. Rokas A, Mead ME, Steenwyk JL, Raja HA, Oberlies NH. Biosynthetic gene clusters and the evolution of fungal chemodiversity. Nat Prod Rep. 2020;37: 868–878.

59. El-Elimat T, Figueroa M, Ehrmann BM, Cech NB, Pearce CJ, Oberlies NH. High-resolution MS, MS/MS, and UV database of fungal secondary metabolites as a dereplication protocol for bioactive natural products. J Nat Prod. 2013;76: 1709–1716.

60. Paguigan ND, El-Elimat T, Kao D, Raja HA, Pearce CJ, Oberlies NH. Enhanced dereplication of fungal cultures via use of mass defect filtering. J Antibiot. 2017;70: 553–561.

61. Li S-M. Genome mining and biosynthesis of fumitremorgin-type alkaloids in ascomycetes. J Antibiot (Tokyo). 2011;64: 45–49.

62. Grundmann A, Kuznetsova T, Afiyatullov SS, Li S-M. FtmPT2, an N-prenyltransferase from Aspergillus fumigatus, catalyses the last step in the biosynthesis of fumitremorgin B. Chembiochem. 2008;9: 2059–2063.

63. Dolan SK, O’Keeffe G, Jones GW, Doyle S. Resistance is not futile: gliotoxin biosynthesis, functionality and utility. Trends Microbiol. 2015;23: 419–428.

64. de Mattos-Shipley KMJ, Simpson TJ. The “emodin family” of fungal natural products-amalgamating a century of research with recent genomics-based advances. Nat Prod Rep. 2023;40: 174–201.

65. Gauthier T, Wang X, Sifuentes Dos Santos J, Fysikopoulos A, Tadrist S, Canlet C, et al. Trypacidin, a spore-borne toxin from Aspergillus fumigatus, is cytotoxic to lung cells. PLoS One. 2012;7: e29906.

66. Mattern DJ, Schoeler H, Weber J, Novohradská S, Kraibooj K, Dahse H-M, et al. Identification of the antiphagocytic trypacidin gene cluster in the human-pathogenic fungus Aspergillus fumigatus. Applied Microbiology and Biotechnology. 2015;99: 10151–10161.

67. Zelder O. [Postoperative serum enzyme changes following abdominal surgery]. Chirurg. 1970;41: 278–280.

68. Throckmorton K, Lim FY, Kontoyiannis DP, Zheng W, Keller NP. Redundant synthesis of a conidial polyketide by two distinct secondary metabolite clusters in Aspergillus fumigatus: Redundant conidial polyketides in*A. fumigatus*. Environ Microbiol. 2016;18: 246–259.

69. Hagiwara D, Takahashi H, Takagi H, Watanabe A, Kamei K. Heterogeneity in Pathogenicity-related Properties and Stress Tolerance in Aspergillus fumigatus Clinical Isolates. Med Mycol J. 2018;59: E63–E70.

70. Turner WB. 1232. The production of trypacidin and monomethylsulochrin by Aspergillus fumigatus. J Chem Soc. 1965; 6658–6659.

71. Russo A, Morrone HL, Rotundo S, Trecarichi EM, Torti C. Cytokine profile of invasive pulmonary aspergillosis in severe COVID-19 and possible therapeutic targets. Diagnostics (Basel). 2022;12: 1364.

72. Gow NAR, Netea MG. Medical mycology and fungal immunology: new research perspectives addressing a major world health challenge. Philos Trans R Soc Lond B Biol Sci. 2016;371. doi:10.1098/rstb.2015.0462

73. Gibbons JG, D’Avino P, Zhao S, Cox GW, Rinker DC, Fortwendel JR, et al. Comparative genomics reveals a single nucleotide deletion in pksP that results in white-spore phenotype in natural variants of Aspergillus fumigatus. Front Fungal Biol. 2022;3: 897954.

74. Dagenais TRT, Keller NP. Pathogenesis of Aspergillus fumigatus in invasive aspergillosis. Clin Microbiol Rev. 2009;22: 447–465.

75. Kim K-H, Willger SD, Park S-W, Puttikamonkul S, Grahl N, Cho Y, et al. TmpL, a transmembrane protein required for intracellular redox homeostasis and virulence in a plant and an animal fungal pathogen. PLoS Pathog. 2009;5: e1000653.

76. Riedling OL, David KT, Rokas A. Global patterns of diversity and distribution in Aspergillus fungi are driven by human and environmental influences. Curr Biol. 2025;35: 4453–4466.e3.

77. Steenwyk JL, Balamurugan C, Raja HA, Gonçalves C, Li N, Martin F, et al. Phylogenomics reveals extensive misidentification of fungal strains from the genus. Microbiol Spectr. 2024;12: e0398023.

78. Price MN, Dehal PS, Arkin AP. FastTree 2--approximately maximum-likelihood trees for large alignments. PLoS One. 2010;5: e9490.

79. Auwera G, Van de A, O’Connor BD. Genomics in the Cloud: Using Docker, GATK, and WDL in Terra. Beijing Boston Farnham Sebastopol Tokyo: O’Reilly; 2020.

80. Poplin R, Ruano-Rubio V, DePristo M, Fennell T, Carneiro M, Van der Auwera G, et al. Scaling accurate genetic variant discovery to tens of thousands of samples. bioRxiv. bioRxiv; 2017. p. 201178. doi:10.1101/201178

81. Wang J. Fast and accurate population admixture inference from genotype data from a few microsatellites to millions of SNPs. Heredity (Edinb). 2022;129: 79–92.

82. Menzel P, Ng KL, Krogh A. Fast and sensitive taxonomic classification for metagenomics with Kaiju. Nat Commun. 2016;7: 11257.

83. Kolmogorov M, Yuan J, Lin Y, Pevzner PA. Assembly of long, error-prone reads using repeat graphs. Nat Biotechnol. 2019;37: 540–546.

84. Walker BJ, Abeel T, Shea T, Priest M, Abouelliel A, Sakthikumar S, et al. Pilon: an integrated tool for comprehensive microbial variant detection and genome assembly improvement. PLoS One. 2014;9: e112963.

85. Simão FA, Waterhouse RM, Ioannidis P, Kriventseva EV, Zdobnov EM. BUSCO: assessing genome assembly and annotation completeness with single-copy orthologs. Bioinformatics. 2015;31: 3210–3212.

86. Gurevich A, Saveliev V, Vyahhi N, Tesler G. QUAST: quality assessment tool for genome assemblies. Bioinformatics. 2013;29: 1072–1075.

87. Minh BQ, Schmidt HA, Chernomor O, Schrempf D, Woodhams MD, von Haeseler A, et al. IQ-TREE 2: New Models and Efficient Methods for Phylogenetic Inference in the Genomic Era. Mol Biol Evol. 2020;37: 1530–1534.

88. Sheikhizadeh S, Schranz ME, Akdel M, de Ridder D, Smit S. PanTools: representation, storage and exploration of pan-genomic data. Bioinformatics. 2016;32: i487–i493.

89. Jonkheer EM, van Workum D-JM, Sheikhizadeh Anari S, Brankovics B, de Haan JR, Berke L, et al. PanTools v3: functional annotation, classification and phylogenomics. Bioinformatics. 2022;38: 4403–4405.

90. Stajich JE, Harris T, Brunk BP, Brestelli J, Fischer S, Harb OS, et al. FungiDB: an integrated functional genomics database for fungi. Nucleic Acids Res. 2012;40: D675–81.

91. Buchfink B, Reuter K, Drost H-G. Sensitive protein alignments at tree-of-life scale using DIAMOND. Nat Methods. 2021;18: 366–368.

92. Terlouw BR, Blin K, Navarro-Muñoz JC, Avalon NE, Chevrette MG, Egbert S, et al. MIBiG 3.0: a community-driven effort to annotate experimentally validated biosynthetic gene clusters. Nucleic Acids Res. 2023;51: D603–D610.

93. Blin K, Shaw S, Kloosterman AM, Charlop-Powers Z, van Wezel GP, Medema MH, et al. antiSMASH 6.0: improving cluster detection and comparison capabilities. Nucleic Acids Res. 2021;49: W29–W35.

94. Tisch D, Schmoll M. Light regulation of metabolic pathways in fungi. Appl Microbiol Biotechnol. 2010;85: 1259–1277.

95. Al Subeh ZY, Raja HA, Monro S, Flores-Bocanegra L, El-Elimat T, Pearce CJ, et al. Enhanced Production and Anticancer Properties of Photoactivated Perylenequinones. J Nat Prod. 2020;83: 2490–2500.

96. Hatmaker EA, Rangel-Grimaldo M, Raja HA, Pourhadi H, Knowles SL, Fuller K, et al. Genomic and phenotypic trait variation of the opportunistic human pathogen Aspergillus flavus and its close relatives. Microbiol Spectr. 2022;10: e0306922.

97. Marim FM, Silveira TN, Lima DS Jr, Zamboni DS. A method for generation of bone marrow-derived macrophages from cryopreserved mouse bone marrow cells. PLoS One. 2010;5: e15263.

98. Winkelströter LK, De Martinis ECP. Different methods to quantify Listeria monocytogenes biofilms cells showed different profile in their viability. Braz J Microbiol. 2015;46: 231–235.

