## Supplementary Tables and Figures for "Opportunistic pathogenicity in fungi can transcend species boundaries"

**Table S1 Secondary metabolites exclusive to *A. fumigatus* relative to *A. fischeri* at 30°C and 37°C.** All compounds detected in at least one *A. fumigatus* strain that were never detected in any *A. fischeri* strain.

| Compound | Condition |
|---|---|
| (-)-Sclerosporin | 30°C and 37°C |
| 10--20-dehydro[12--13-dehydroprolyl-2-(1--1-dimethylallyl)tryptophyl]diketopiperazine] | 30°C and 37°C |
| 12--13-Dehydroprolyl-2-(1--1-dimethylallyltryptophyl)diketopiperazine | 30°C and 37°C |
| 12--13-Dihydroxyfumitremorgin C | 30°C and 37°C |
| 19--20-Epoxycytochalasin D | 30°C only |
| 1-hydroxyisorhodoptilometrin | 30°C only |
| 2-Hydroxyacoronene | 30°C and 37°C |
| 5--8-Epidioxyergosta-6--9(11)--22-trien-3-ol | 30°C and 37°C |
| 5-hydroxy-7-methoxy-4-methylcoumarin | 30°C and 37°C |
| Asterriquinone-analogue | 30°C and 37°C |
| Bisdechlorogeodin | 30°C and 37°C |
| C-11-epimer-verruculogen-TR-2 | 30°C and 37°C |
| Crotocin | 30°C and 37°C |
| Cyclo (Pro-Phe) | 30°C and 37°C |
| Cyclo(L-Pro-L-Leu)/Cyclo(S-Pro-S-Leu) | 30°C and 37°C |

| | |
|---|---|
| Desmethyl dermoquinone | 30°C and 37°C |
| Diacetoxyscirpenol | 30°C only |
| Diepoxin (zeta) | 30°C and 37°C |
| Ergosterol peroxide | 30°C and 37°C |
| Fumagillin | 30°C and 37°C |
| Fumigaclavine C | 30°C and 37°C |
| Fusaproliferin | 30°C and 37°C |
| Isoherqueinone/Herqueinone | 30°C only |
| Meleagrin | 30°C only |
| Methyl-asterrate | 30°C and 37°C |
| Monomethylsulochrin | 30°C and 37°C |
| New_89 | 30°C and 37°C |
| Phomochromenone C | 30°C and 37°C |
| Pseudaboydin B | 30°C and 37°C |
| Pseurotin | 30°C and 37°C |
| Questin | 30°C and 37°C |
| Ramiferin | 30°C and 37°C |
| RFJ86-I | 30°C and 37°C |

| | |
|---|---|
| Rhizoctonic acid | 30°C only |
| Soudanone H | 30°C and 37°C |
| S-Sydonol | 30°C and 37°C |
| Sulochrin | 30°C only |
| Thielavin_Z8 | 30°C and 37°C |
| Trichodermol | 30°C and 37°C |
| Trichothecin | 30°C and 37°C |
| Trichothecinol D | 30°C only |
| Trypacidin | 30°C and 37°C |
| Zearalanone | 30°C and 37°C |
| Zearalenol-(alpha)/Zearalenol-(beta) | 30°C and 37°C |

**Table S2 Phenotype differences between species**

Results of one-way ANOVAs testing for species-level differences across all measured traits, reporting F-statistics, effect sizes ($\eta^2$), and Benjamini–Hochberg (FDR)-adjusted p-values, with traits ordered by the proportion of variance explained by species.

| Trait | F statistic | $\eta^2$ | p-value | BH padj |
|---|---|---|---|---|
| Radial Growth CFW 25 µg/mL | 80.42851 | 0.715375 | 3.03E-10 | 5.21E-09 |
| Biofilm 4 °C | 77.0687 | 0.706607 | 4.96E-10 | 5.21E-09 |
| Radial Growth MM 44 °C | 37.32733 | 0.538422 | 7.91E-07 | 5.53E-06 |
| Radial Growth YAG 44 °C | 33.46863 | 0.511216 | 2.02E-06 | 1.06E-05 |
| Radial Growth MM 30 °C | 25.71028 | 0.445506 | 1.62E-05 | 6.80E-05 |
| Biofilm 37 °C | 24.5923 | 0.434552 | 2.24E-05 | 6.82E-05 |
| Radial Growth YAG 30 °C | 24.53961 | 0.434025 | 2.27E-05 | 6.82E-05 |
| CFW 37 °C | 20.2805 | 0.387917 | 8.36E-05 | 2.19E-04 |
| WGA 4 °C | 17.86458 | 0.358262 | 1.85E-04 | 4.31E-04 |
| Radial Growth YAG 37 °C | 14.3911 | 0.310213 | 6.23E-04 | 1.31E-03 |
| Radial Growth CR 50 µg/mL | 12.76776 | 0.2852 | 1.14E-03 | 2.18E-03 |
| CFW 4 °C | 4.819174 | 0.130888 | 3.55E-02 | 6.21E-02 |
| Radial Growth MM 37 °C | 4.510998 | 0.123552 | 4.15E-02 | 6.70E-02 |
| WGA 37 °C | 4.125849 | 0.114208 | 5.06E-02 | 7.51E-02 |
| Radial Growth 10% BFS | 4.013443 | 0.111443 | 5.37E-02 | 7.51E-02 |

| | | | | |
|---|---|---|---|---|
| Radial Growth T-butyl | 3.732474 | 0.104456 | 6.23E-02 | 8.17E-02 |
| Radial Growth Fe starvation | 3.047309 | 0.086948 | 9.05E-02 | 1.12E-01 |
| Radial Growth 0.02 mM menadione | 0.776678 | 0.023696 | 3.85E-01 | 4.49E-01 |
| Dectin 4 °C | 0.488969 | 0.01505 | 4.89E-01 | 5.41E-01 |
| Adhesion | 0.056664 | 0.001768 | 8.13E-01 | 8.54E-01 |

**FIGURE S1:** ***A fumigatus*** **SNV tree and population genomics**

Single nucleotide variant (SNV) tree built (FastTree) from 7,386 pruned and thinned SNVs shared among 408 clinical and environment strains. Of these, 322 received population assignments (consensus between ADMIXTURE and DAPC) and are color coded by population.

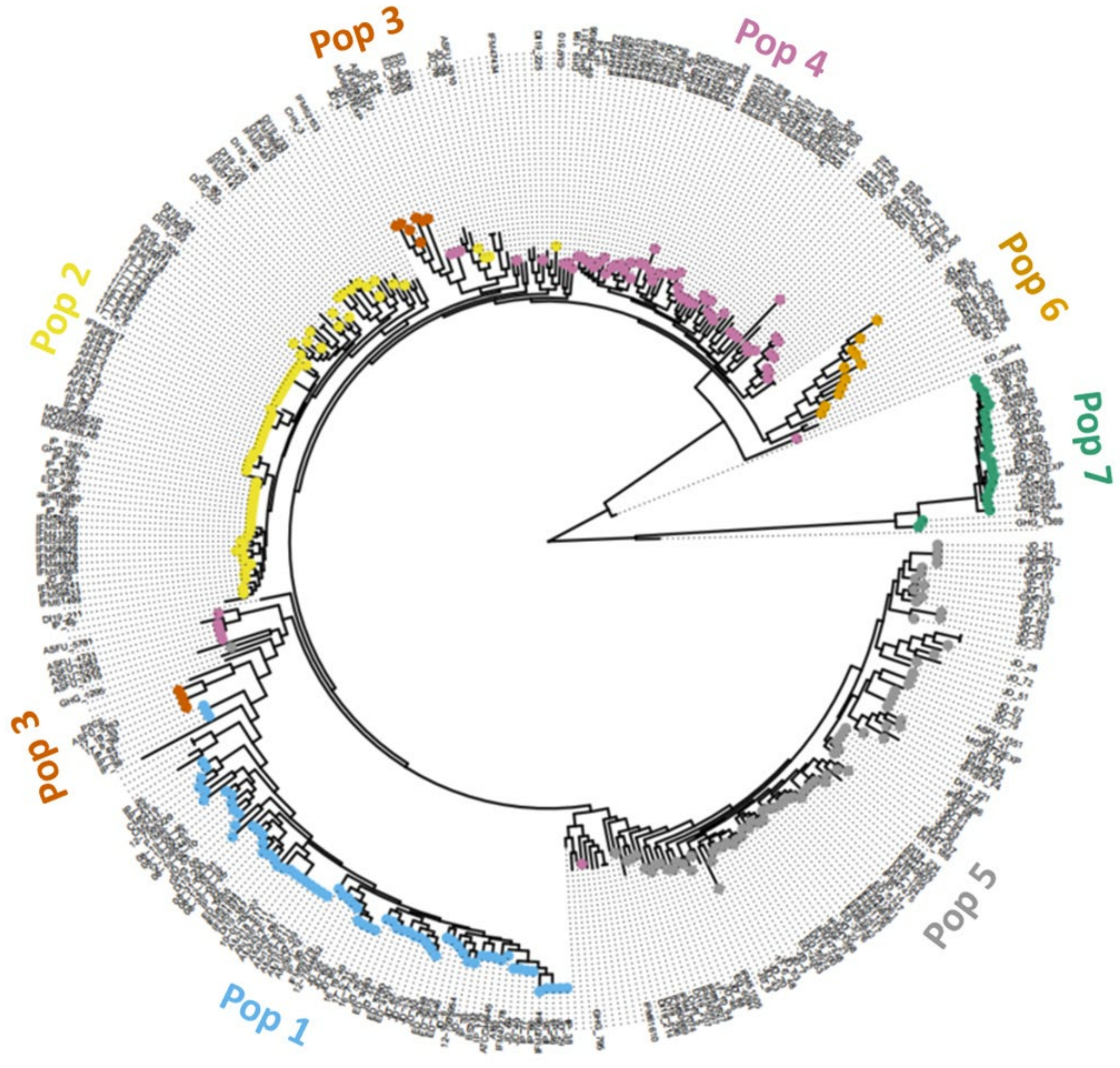

**FIGURE S2: *A. fumigatus* pangenome content**

Rarefaction curves for pangenomes of the 16 *A. fumigatus* strains. Points show gene counts from the random subsampling (without replacement) of the genomes of 16 *A. fumigatus* strains. Subsample size was incrementally increased from one to 16 genomes, with 10,000 replicates performed at each step. Results illustrate that the *A. fumigatus* pangenome is open (Heaps' law regression, $\gamma = 0.854$).

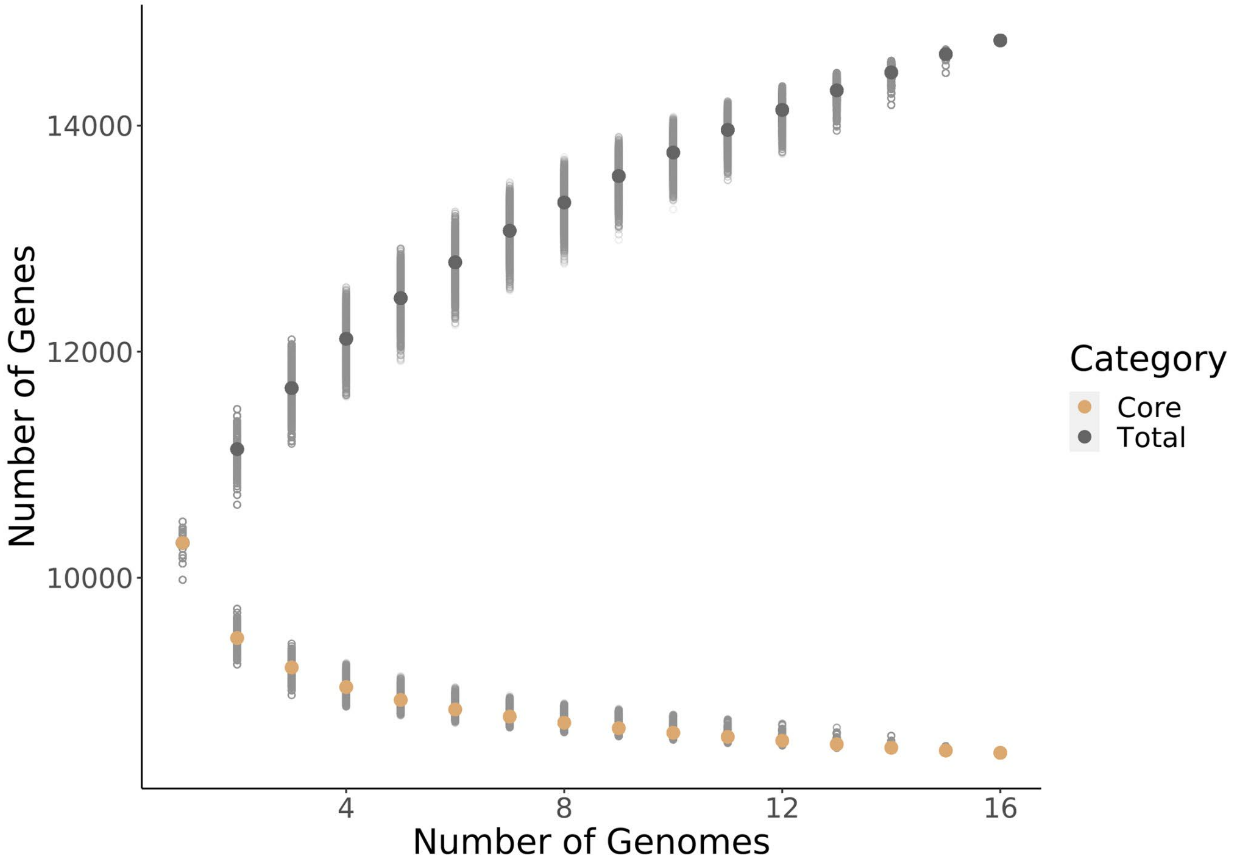

**Figure S3: *A. fumigatus* and *A. fischeri* molecular phylogeny**

Maximum likelihood phylogeny of 3,135 BUSCOS (Methods). Distances reported and highlighted on the tree are the median, pairwise patristic distances between all *A.fumigatus* and *A. fischeri* strains (purple) and then all focal strains relative to the outgroup (green). Inset shows patristic distances between all strains of each species.

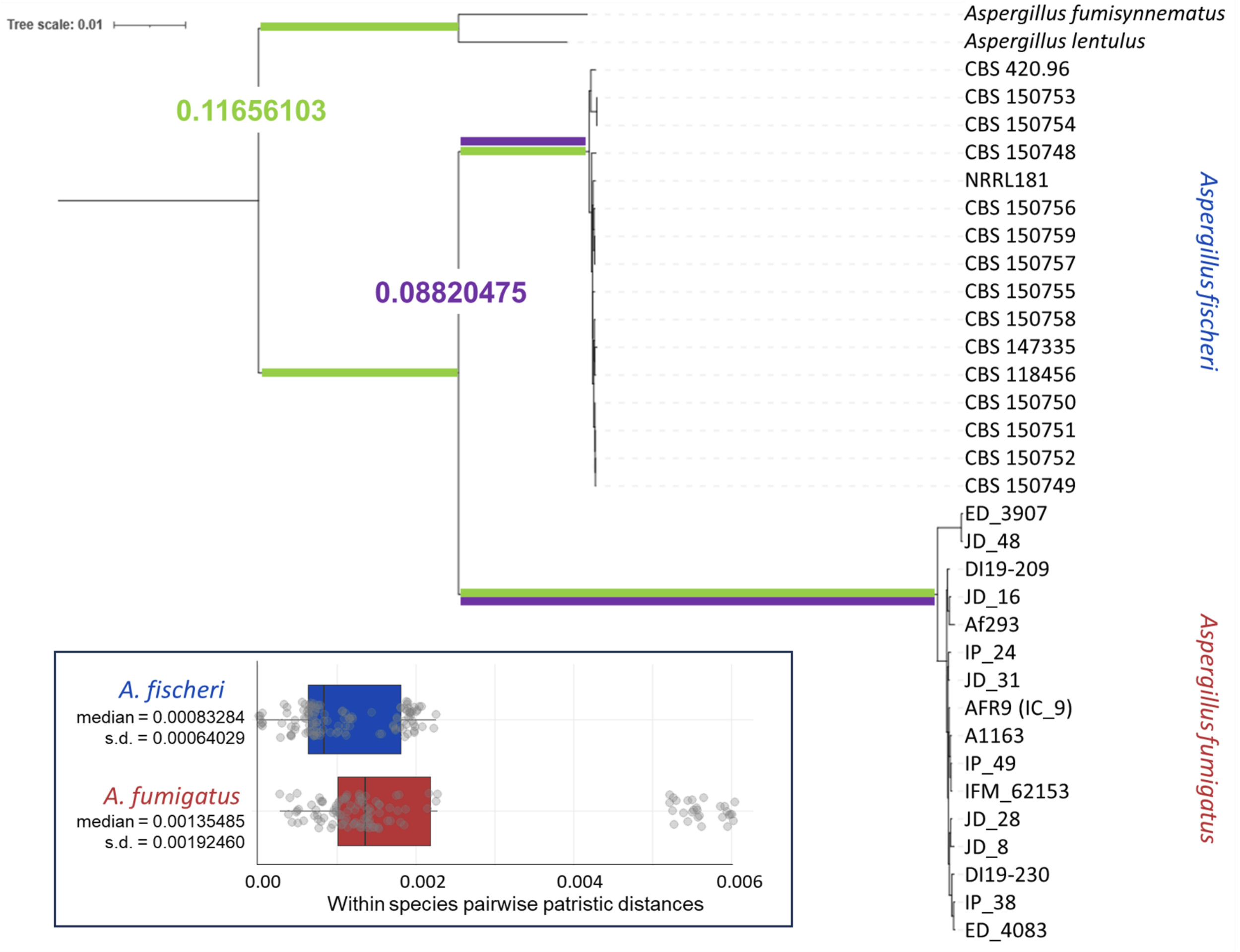

**FIGURE S4: In vitro growth and cell wall phenotype loadings**

PCA results and loadings for all monoculture phenotypes assayed in all strains of *A. fumigatus* (red points) and *A. fischeri* (blue points). Abbreviations are identical to those in Figure 1. MM=minimal media; YAG=yeast extract glucose media; CFW=Calcofluor White; WGA=Wheat Germ Agglutinin); dectin (solubilized pattern recognition receptor specific to β-(1,3)-glucans); menadione (inducer of oxidative stress); CW=Congo red (inducer of oxidative stress).

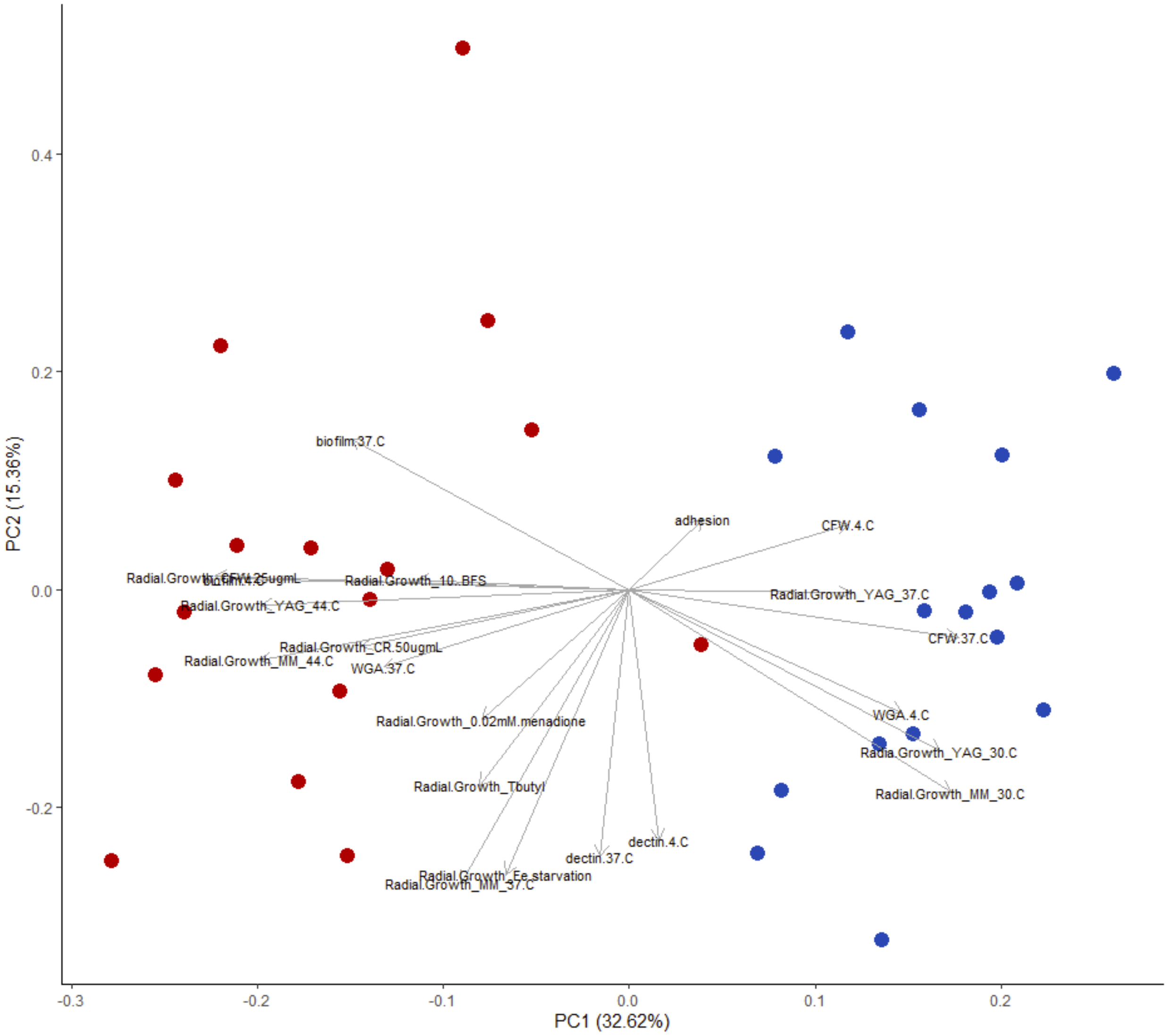

**FIGURE S5: Tanglegram comparing all monoculture phenotypes to strain/species tree.**

The BUSCO gene tree (left) compared to the phenetic tree (right). The phenetic tree is a neighbor-joining tree based on phenotypic distances (Euclidean distances among z-scores). Metrics comparing the tree topologies are as follows:

- Normalized RF distance (0=identical, 1=completely different): 0.8571
- Normalized quartet distance (0=identical, 1=completely different):0.3179
- Correlation of patristic distances (molecular matrix and z scores) : 0.609

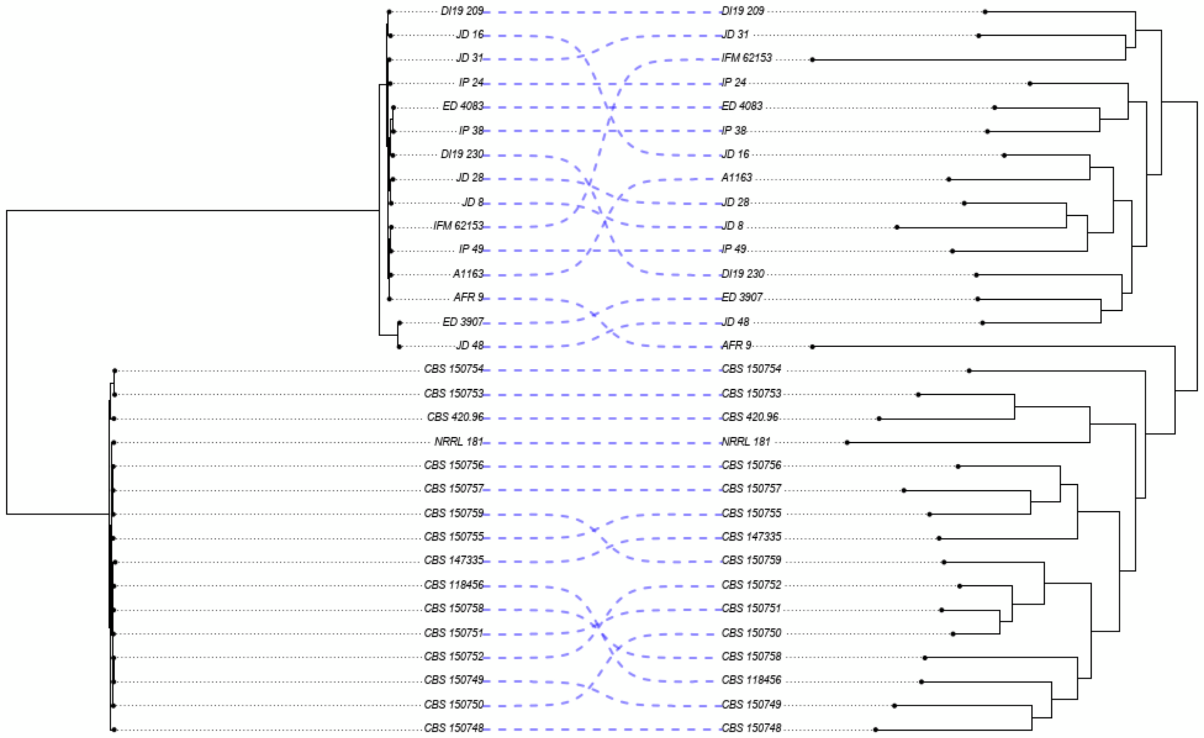

**FIGURE S6: Count of metabolites detected in each species at two temperatures.**

Summary of unique compounds detected across strains of both species under two different growth (temperature) conditions.

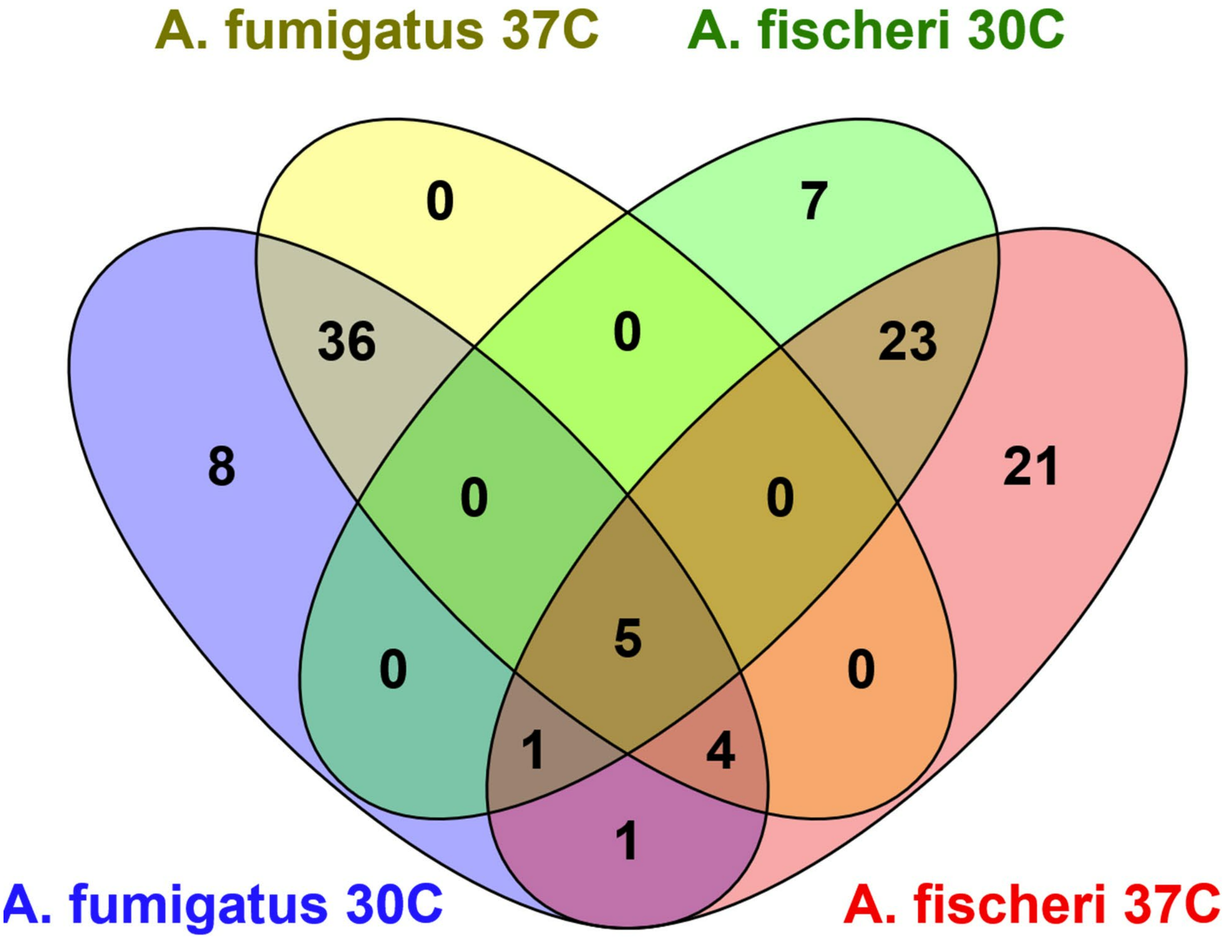

**FIGURE S7: Metabolite profiling**

Principal component analysis based on retention times and individual peak areas for different metabolites from each of the *A. fumigatus* strains following growth at either 30 °C or 37 °C. Each point is derived from the average features of three biological replicates. Arrows highlight the shift in PCA space of the two strains showing the greatest, temperature-dependent changes in chemical profiles.

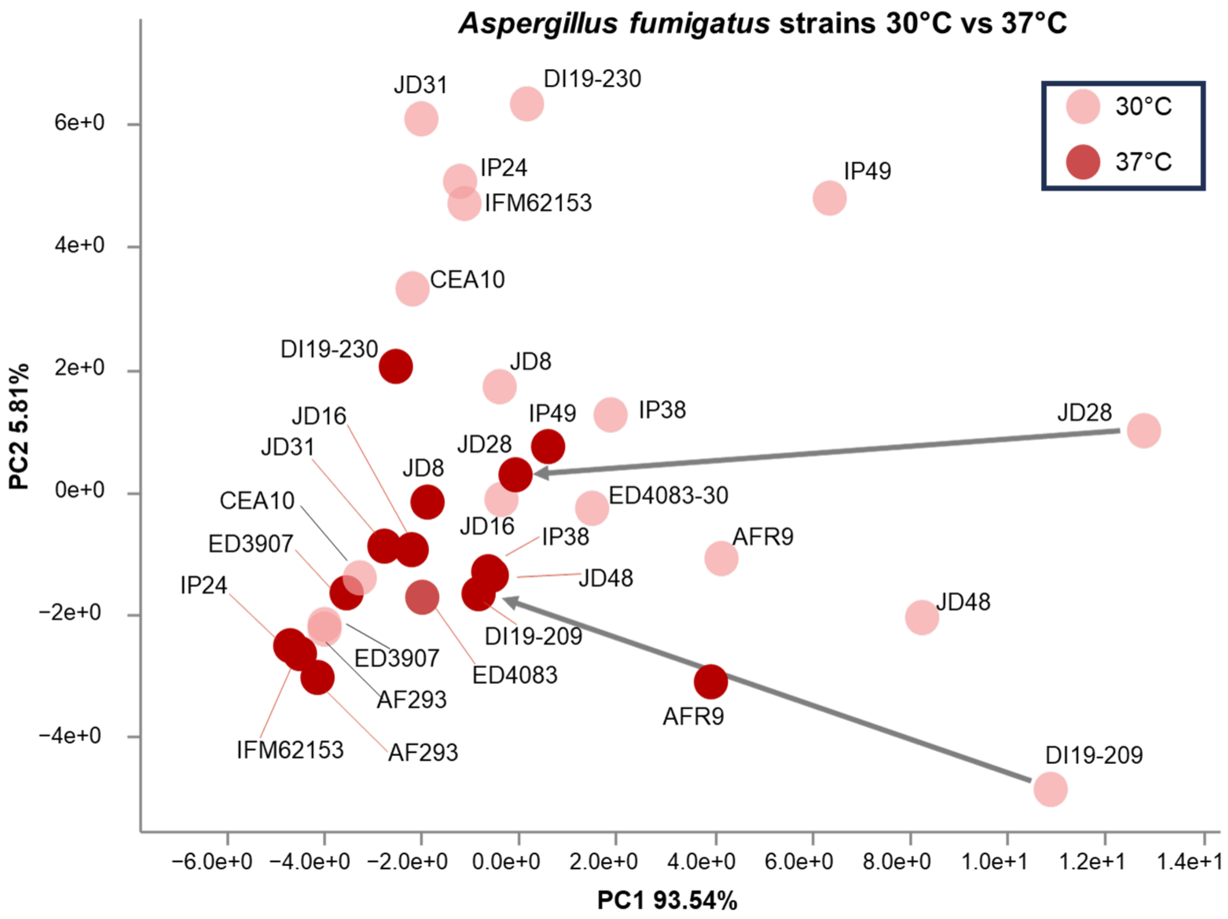

**FIGURE S8: Correlations of responses in macrophages**

Spearman correlations of all *in vivo* responses to macrophages among all strains of: *A. fischeri* (left), *A. fumigatus* (middle), and both species (right).

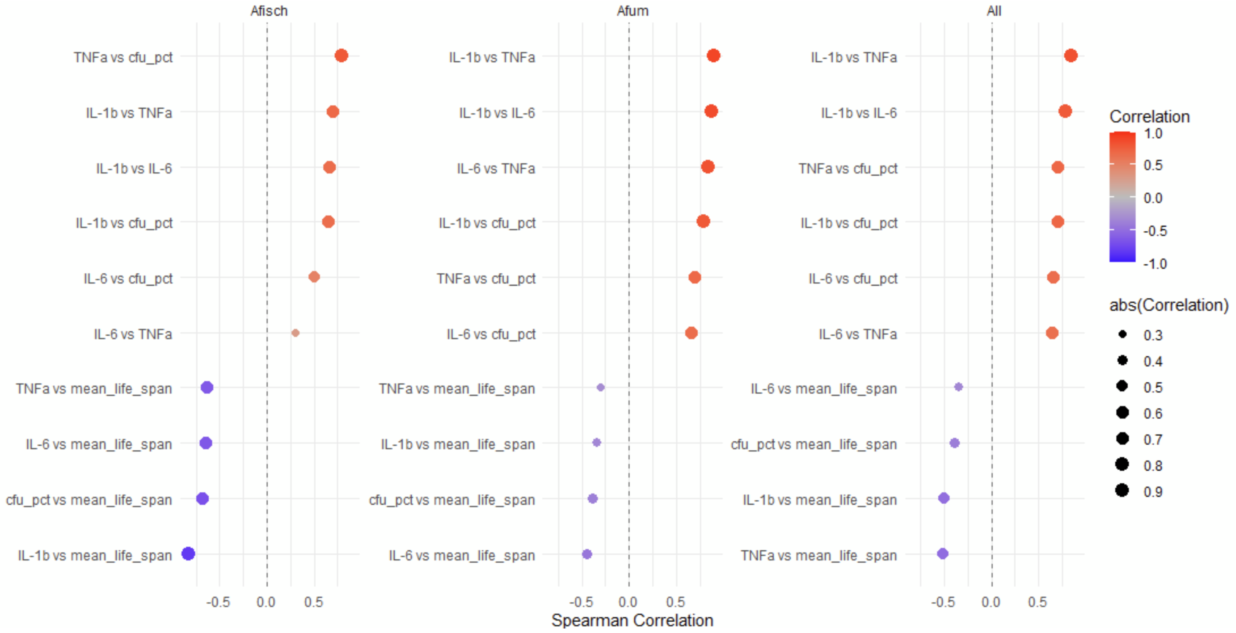

**FIGURE S9: Correlation between *in vitro* macrophage response to spores and lifespan of infected mice**

Spearman correlation among all strains between conidial survival and mean (n=20 for each strain) lifespan over the 15 days post infection.

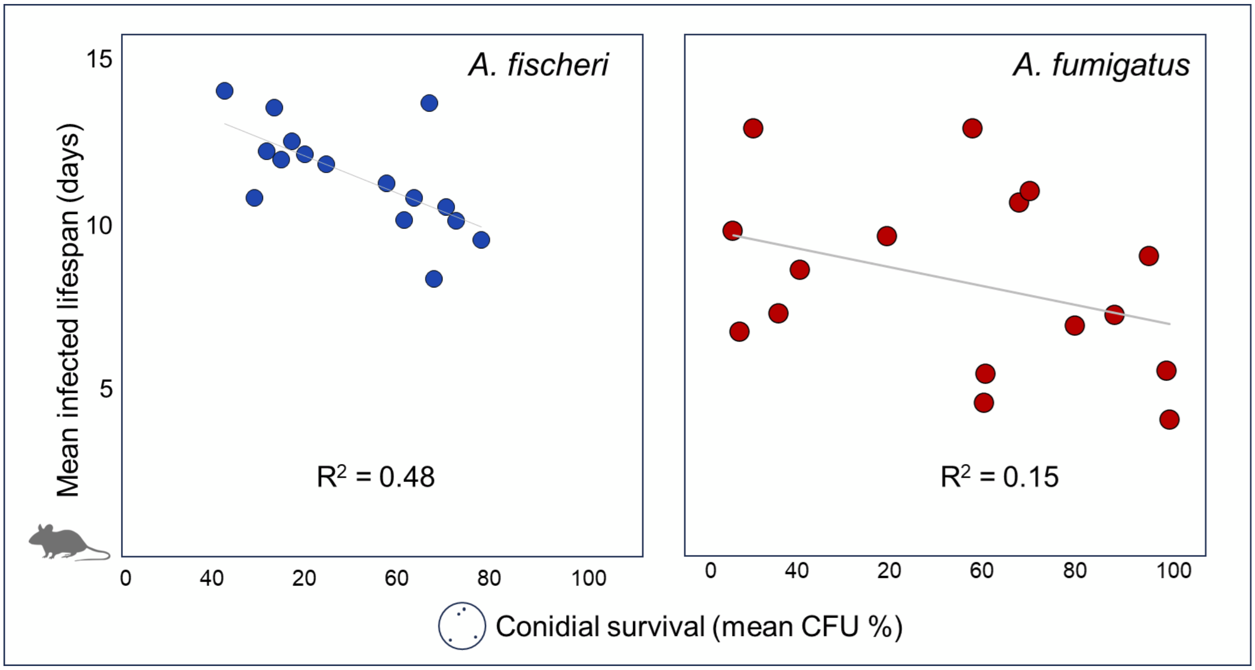

**FIGURE S10: PCA of *in vitro* macrophage response to spores**

Shown are the PCA results and loadings for all measured *in vivo* responses to macrophages. Strains of *A. fumigatus* (red points) and *A. fischeri* (blue points) are selectively labeled to identify strains discussed along with the reference strains NRRL 181 and CEA10(A1163).

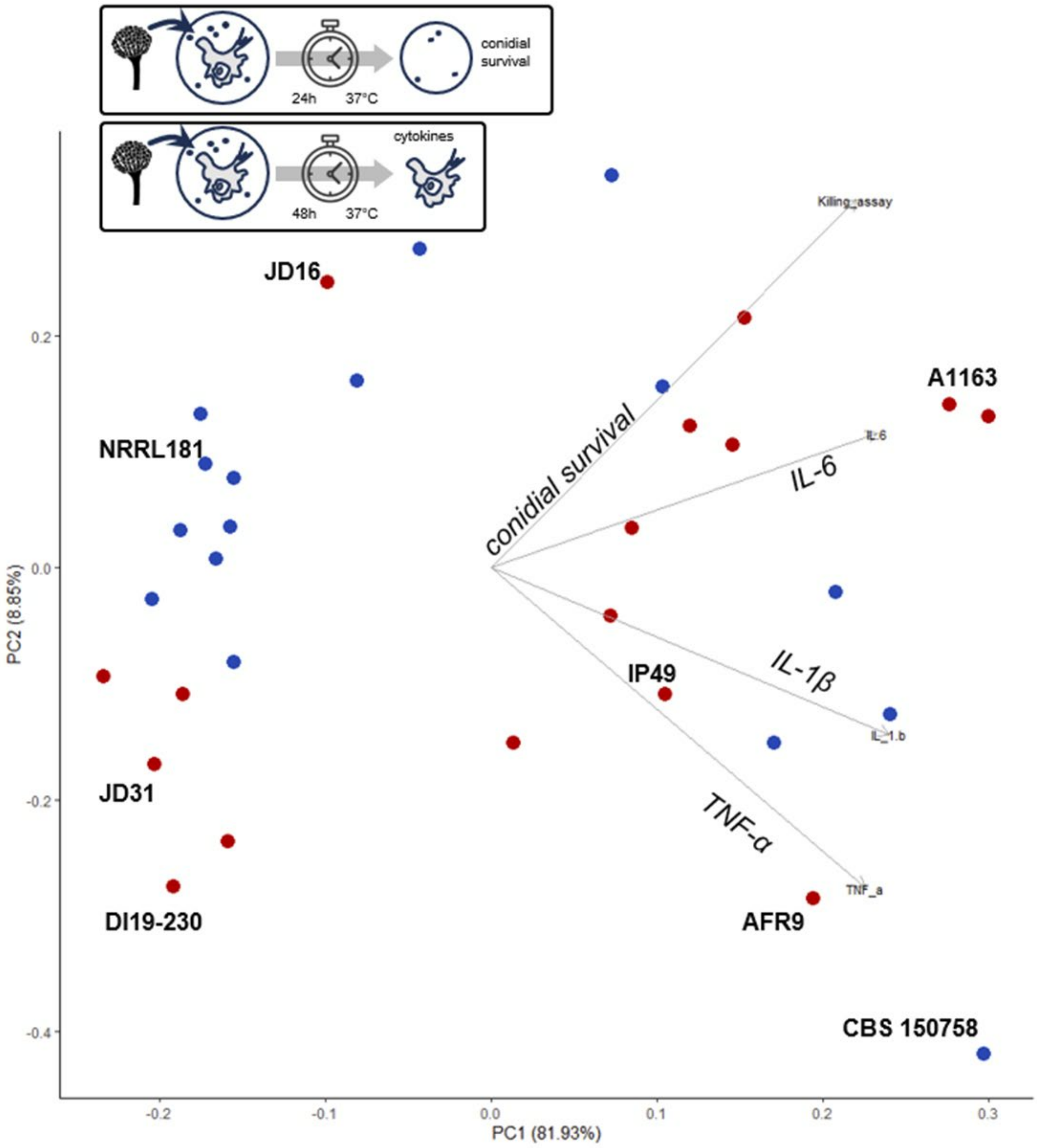

**FIGURE S11: PCA of combined *in vitro* and *in vivo* results.**

PCA results and loadings for all measured *in vivo* responses to macrophages and virulence results from an *in vitro* murine model of pulmonary aspergillosis.

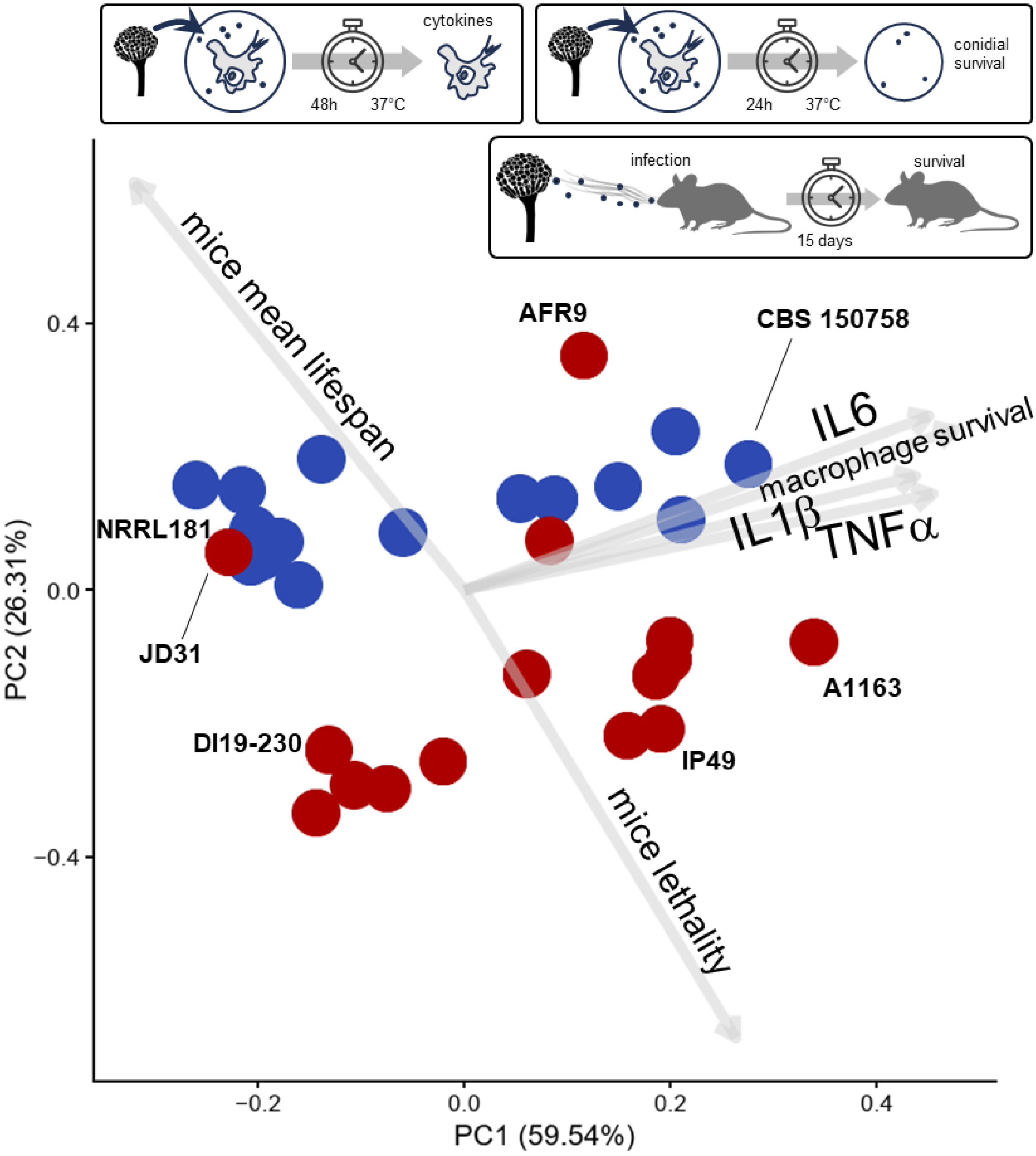

**Figure S12: Tanglegram of *in vivo* and *in vitro* phenotypes**

The BUSCO gene tree (left) compared to the phenetic tree of *in vivo* and *in vitro* phenotypes (right). The phenetic tree is a neighbor-joining tree based on phenotypic distances (Euclidean distances among z-scores). Metrics comparing the tree topologies are as follows:

- Normalized RF distance (0=identical, 1=completely different): 0.9643
- Normalized quartet distance (0=identical, 1=completely different):0.4119
- Correlation of patristic distances: 0.2507

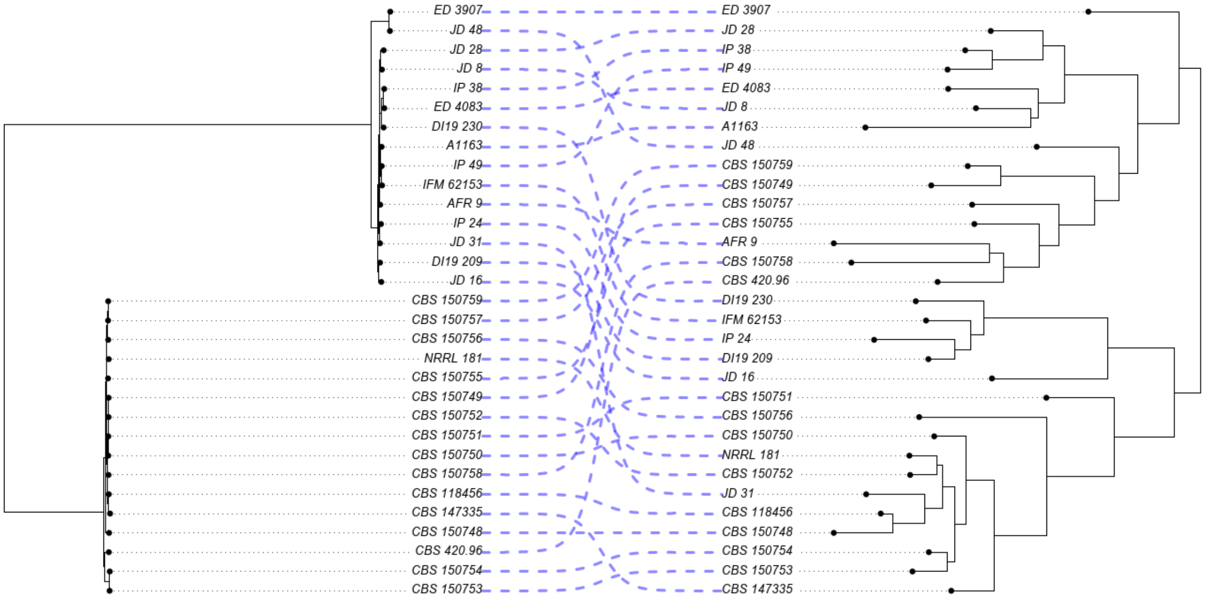

**Figure S13: All monoculture, *in vivo*, and *in vitro* phenotypes**
Heatmap of all assayed traits, crusted by z-score.

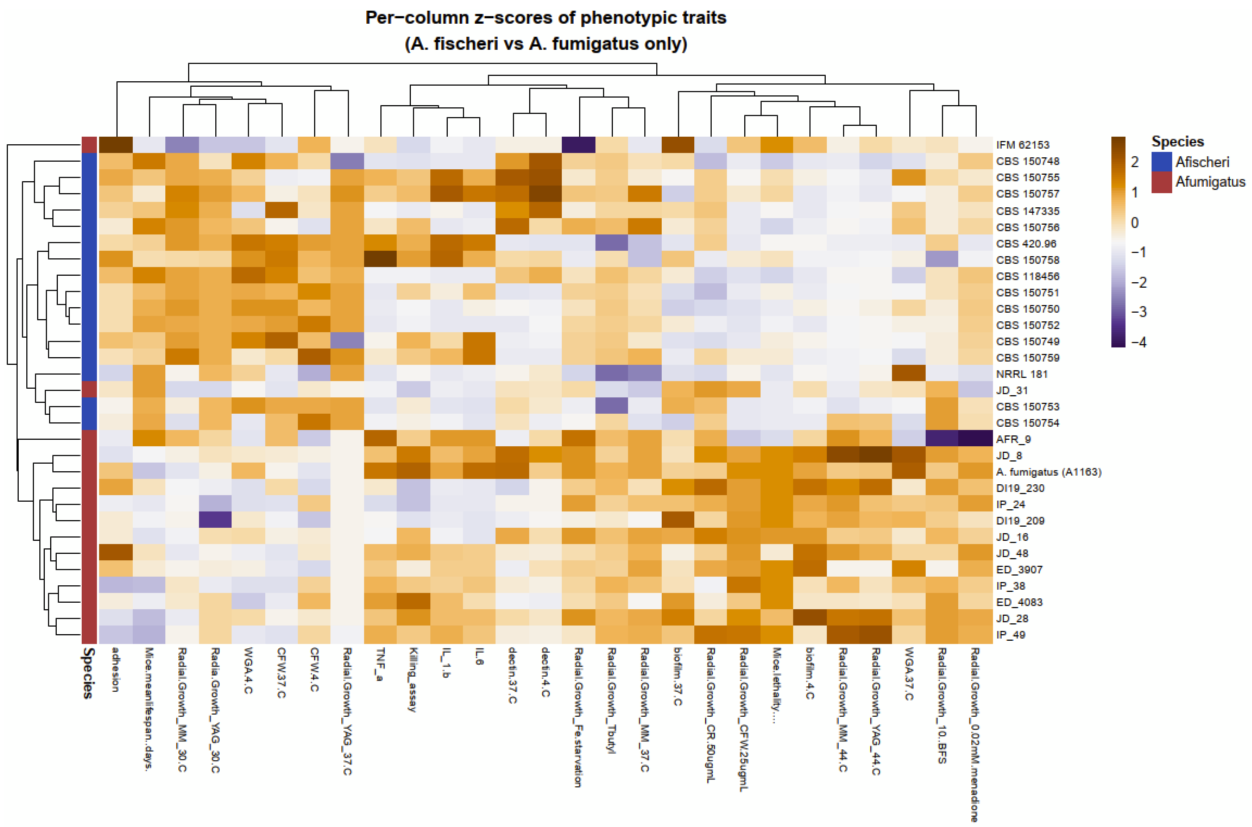
